# Transmission of Cervid prions to Humanized Mice Demonstrates the Zoonotic Potential of CWD

**DOI:** 10.1101/2022.04.19.488833

**Authors:** Samia Hannaoui, Irina Zemlyankina, Sheng Chun Chang, Maria Immaculata Arifin, Vincent Béringue, Debbie McKenzie, Hermann M. Schatzl, Sabine Gilch

## Abstract

Prions cause infectious and fatal neurodegenerative diseases in mammals. Chronic wasting disease (CWD), a prion disease of cervids, spreads efficiently among wild and farmed animals. Potential transmission to humans of CWD is a growing concern due to its increasing prevalence. Here, we provide the strongest evidence to date supporting the zoonotic potential of CWD prions, and their probable materialization in humans using mice expressing human prion protein (PrP) as an infection model. Inoculation of these mice with deer CWD isolates resulted in atypical clinical manifestations, with prion seeding activity and efficient transmissible infectivity in the brain and, remarkably, in feces. Intriguingly, the protease-resistant PrP in the brain resembled that found in a familial human prion disease and was transmissible upon second passage. Our results are the first evidence that CWD can infect humans with a distinctive clinical presentation, signature, and tropism, and might be transmissible between humans while current diagnostic assays might fail to detect it. These findings have major implications for public health and CWD management.

## Introduction

Prions are the causative agents of transmissible and fatal neurodegenerative diseases of humans (e.g., Creutzfeldt-Jakob disease (CJD)) and animals (scrapie in sheep, bovine spongiform encephalopathy (BSE) in cattle, and CWD in cervids) ^1^. Prion diseases are characterized by the accumulation in the brain of the infectious prion protein, PrP^Sc^, derived after a structural transition from its host-encoded, cellular isoform, PrP^C^ ^2^. CWD is the only prion disease known to affect both free-roaming and farmed animals. Cervid species naturally affected by CWD include white-tailed deer (WTD; *Odocoileus virginianus*), mule deer (*O. hemionus*), elk (*Cervus canadensis*), red deer (*C. elaphus*), moose (*Alces alces* sp.), and reindeer (*Rangifer tarandus tarandus*) ^3^. As of today, it has been identified in 30 U.S. states, with a prevalence as high as 40% in certain endemic areas ^4^, 4 Canadian provinces, South Korea, and 3 Northern European countries, Norway, Finland, and Sweden ^3^. For most prion diseases, with the exception of scrapie and CWD, infectious prions are mostly confined to the central nervous system. In contrast, in cervids affected with CWD, infectivity has been found in the lymphatic system, salivary gland, intestinal tract, muscles, antler velvet, blood, urine, saliva, and feces ^3^, demonstrated to be transmissible ^5^. CWD prions are shed into the environment via bodily fluids and excreta. They bind to soil and are taken up by plants, making the environment infectious for decades to come ^3, 6^. The persistence of CWD prions in the environment amplifies the already effective transmission within and between cervid species. Therefore, CWD is considered to be the most contagious prion disease with fast spreading and efficient horizontal transmission.

Zoonotic BSE (i.e., variant CJD) provides undeniable evidence that animal prions can infect humans, resulting in distinct disease manifestation and strain properties ^7^. Epidemiological studies in CWD endemic areas have neither indicated an increased incidence of CJD patients, nor unusual prion disease subtypes ^4, 8, 9^. Numerous studies assessing the zoonotic potential of CWD, both *in vitro* and *in vivo,* overall conclude that the risk of CWD crossing the human barrier is low ^8, 10–20^. However, prions are dynamic and evolving, and interspecies passage of CWD can result in prion adaptation to new host species. The existence of more than one CWD strain ^3^ may contribute to higher heterogeneity in disease and transmission profiles. *In vitro* studies using protein misfolding cyclic amplification (PMCA) demonstrated that PrP^CWD^ can convert human PrP^C^ into PrP^CWD^, though either after prion strain stabilization and adaptation *in vitro* or *in vivo*, or at low to moderate efficiencies ^18, 21^. Efficiency of PMCA conversion also depended on a human PrP polymorphism at position 129 (methionine (M)/valine (V)), affecting susceptibility to prion disease in humans ^17, 22, 23^, especially in the only to date known human prion disease (vCJD) acquired after animal prions (BSE) crossed the species barrier ^7^. In contrast, most *in vivo* studies indicated that inoculation of different CWD prion isolates into transgenic mice overexpressing human PrP^C^ with different genotypes at codon 129 did not result in disease ^10–12, 14, 15, 19^. A recent study by Wang *et al.*, showed that transgenic mice overexpressing human M129- and V129-PrP^C^ are susceptible to *in vitro*-generated PrP^CWD^, with elk CWD used as a seed and human V129-PrP^C^ used as a substrate in PMCA. In addition, in this study, PrP^CWD^ could only convert V129-PrP^C^ substrate *in vitro*, but not M129-PrP^C^ ^20^.

Squirrel monkeys were found to be susceptible to intracerebral (i.c.) and oral CWD infection ^18, 21, 24^, while *Cynomolgus* macaques have conflicting results. Race and collaborators have reported that the latter non-human primate model was not susceptible to CWD ^16, 25, 26^; while a consortium study including our group reported an atypical phenotype, and positive RT-QuIC and PMCA assays in harvested tissues from macaques after challenge with CWD using different routes of inoculation (Czub S, PC, 2017). Although the human species barrier to CWD infection is presented as strong by most of the studies, yet these findings are still a matter of debate ^3^. In the absence of effective management strategies, the prevalence of CWD, the affected geographical areas and the host species range of CWD are increasing, and with that, the potential for human exposure to CWD prions is also increasing. In addition, infectious prions have been found in skeletal muscle ^27^ and antler velvet ^28^ in infected cervids, raising yet more concerns about zoonotic transmission of CWD through venison consumption and/or application of cervid products in traditional medicines ^3^.

In this study, we evaluated the zoonotic potential of CWD using a transgenic mouse model overexpressing human M129-PrP^C^ (tg650 ^29^). We inoculated tg650 mice intracerebrally with two deer CWD isolates, Wisc-1 and 116AG ^30–33^. We demonstrate that this transgenic line was susceptible to infection with CWD prions and displayed a distinct leading clinical sign, an atypical PrP^Sc^ signature and unusual fecal shedding of infectious prions. Importantly, these prions generated by the human PrP transgenic mice were transmissible upon passage. Our results are the first evidence of a zoonotic risk of CWD when using one of the most common CWD strains, Wisc- 1/CWD1. We demonstrated in a human transgenic mouse model that the species barrier for transmission of CWD to humans is not absolute. The fact that its signature was not typical raises the questions whether CWD would manifest in humans as a subclinical infection, whether it would arise through direct or indirect transmission including an intermediate host, or a silent to uncovered human-to-human transmission, and whether current detection techniques will be sufficient to unveil its presence.

## Results

To assess and understand the zoonotic potential of CWD, we transmitted CWD prions from deer, without prior adaptation through *in vivo* or *in vitro* amplification, to tg650 mice overexpressing homozygously human M129-PrP^C^ ^29^ (humanized mice). We inoculated intracerebrally tg650 mice (n = 10 per group) with brain homogenates of CWD-positive WTD. To investigate potential differences in the ability of CWD strains to infect tg650 mice, two isolates harboring different strains of CWD were used, Wisc-1, a CWD strain derived from a WTD expressing the cervid wild-type PrP, or 116AG, an isolate from a WTD harboring a polymorphism at position 116 (A116G) and identified to contain a mixture of two co-existing strains both distinct from Wisc-1 prions ^30–33^. We also had a group of nine non-infected tg650 age-matched negative controls (623 – 934 days post-inoculation (dpi)). The scheme in Figure S1 summarizes transmissions and results detailed onwards.

### Humanized mice are susceptible to CWD prions

In cervids affected with CWD, the clinical presentation is mainly that of a wasting syndrome. Clinical manifestations such as behavioral changes (e.g., depression, isolation from the herd), excessive salivation, polyuria and teeth grinding are also observed. However, in rodent models of prion disease, including CWD inoculated models, we usually anticipate neurological signs, such as ataxia, gait abnormalities, weakness, rigid tail, and kyphosis, and behavioral changes such as isolation, wasting, irresponsiveness, and weight loss.

We closely monitored tg650 CWD-inoculated mice, and their age-matched non-infected controls, for progressive signs of prion diseases. Starting at about one-year post-inoculation, we observed that 93.75% of the mice, irrespective of the inoculum, developed myoclonus, as diagnosed by our veterinarian, an unusual clinical manifestation not typically observed in rodent prion models. Clinical signs progressed, with some mice undergoing cycles of weight loss and (re)gain, and lastly drastic weight loss. Eventually, some animals developed typical signs of prion disease, such as rigid tail, kyphosis, hind-limb clasping, ataxia, paralysis, heavy breathing, irresponsiveness, and gait abnormalities (Table S1). Mice were euthanized either at a terminal disease stage or at the experimental endpoint of >900 dpi.

Animals with subtle clinical signs were identified as those that developed some subtle and/or transient signs that did not progress over the course of the experiment; however, all of these animals had myoclonus. Animals with terminal clinical signs were those with confirmatory prion signs that progressed and reached terminal stage of disease. Based on this standard, of the mice inoculated with Wisc-1 prions 77.7% were clinical, of which, 44.4% progressed with terminal clinical signs, and 33.3% developed subtle clinical signs (Table S2, and Figure S2). Of the 116AG- inoculated mice, 71.5% developed progressive clinical signs, of these, 28.6% showed terminal clinical signs, and 42.9% presented with subtle clinical signs (Table S2 and Fig S2). Age-matched non-infected control mice (n = 9) did not exhibit any behavioural or neurological signs and were healthy up to the experimental endpoints between 623 dpi and >900 dpi (Table S2, and Figure S2). We used real-time quaking-induced conversion (RT-QuIC) assay, a highly sensitive *in vitro* conversion technique demonstrated to vie the sensitivity of animal bioassays ^34^, to detect the presence of PrP amyloid seeding activity in the brains, spinal cords (SC), and spleens of CWD-inoculated tg650 mice.

The RT-QuIC results depicted in Figure 1A show reactions seeded with brain homogenates and are representative of one assay for each tg650-inoculated mouse with Wisc-1. Each curve represents the average of 8 replicates for each tested dilution (10^-1^ to 10^-6^). In addition, to achieve a better sensitivity of detection, we included a substrate replacement step after 25 hours reaction time (arrow shown in Figure 1B), where 90% of the reaction mixture was replaced with fresh assay buffer containing recombinant PrP and the RT-QuIC run was carried on up to 60 hours. Brain homogenates of age-matched non-infected control mice were also tested. For each RT-QuIC run, an age-matched negative control BH was used as internal negative control, and they were consistently negative (Figure 1B). WTD Wisc-1 (Figure 1B) and sCJD brain homogenates (Figure S3A) were used as positive controls.

**Figure 1.**
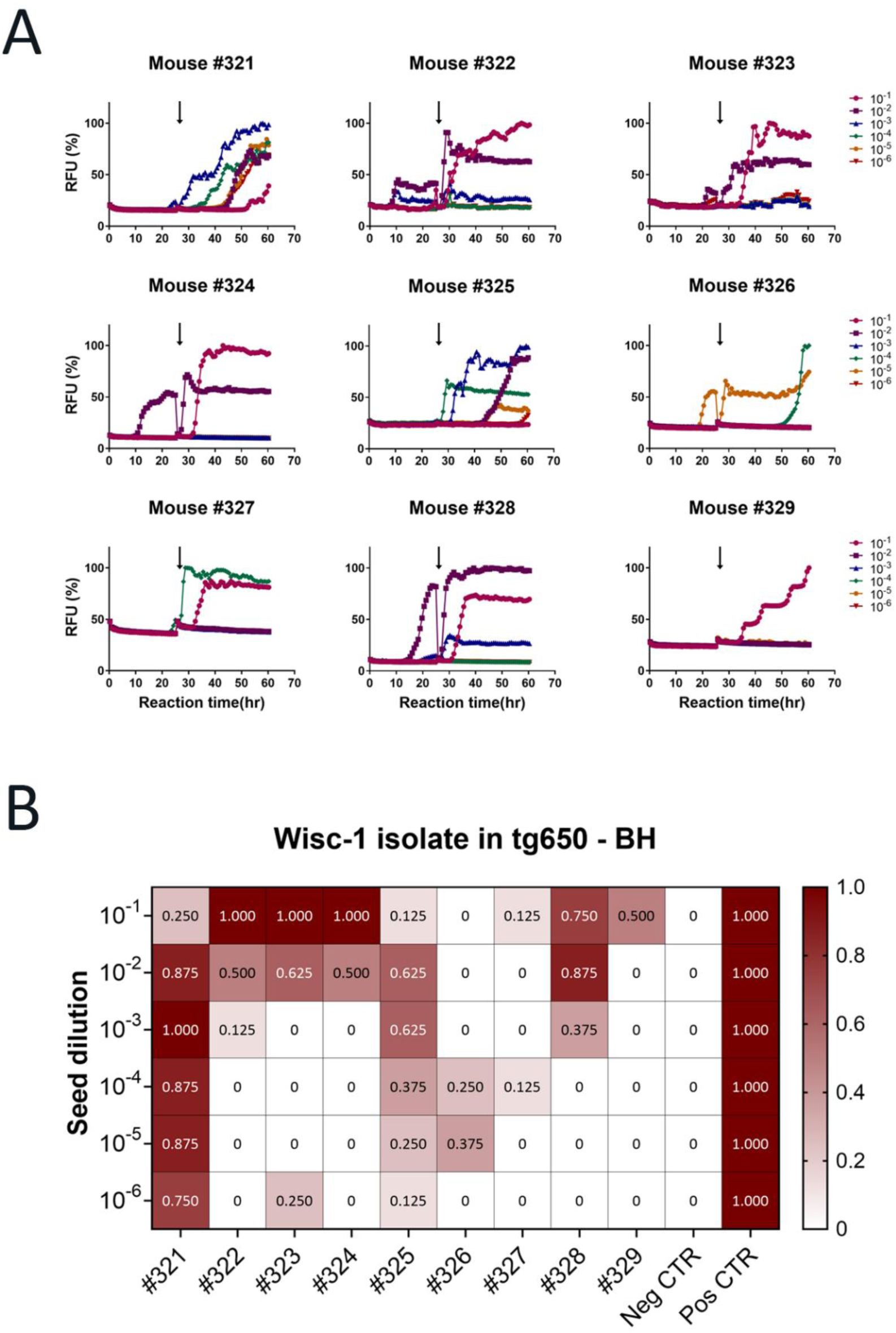
Prion seeding activity in brains of CWD-Wisc1-inoculated transgenic mice overexpressing human PrP (129MM). Transgenic tg650 mice were intracerebrally inoculated with 1% WTD-Wisc-1 isolate. **A.** The graphs depict representative RT-QuIC results of serially diluted (10^-1^ to 10^-6^) tg650 brain homogenates using mouse rPrP substrate. Replacement with fresh assay buffer containing rPrP is indicated with a black arrow. Fluorescence signals were measured every 15 minutes. The x-axis represents the reaction time (hours), the y-axis represents the relative fluorescence units, and each curve represents a different dilution. Mean values of eight replicates were used for each dilution. The cut-off was based on the average fluorescence values of negative control +5SD used in every assay. **B.** Summary of the RT-QuIC analysis of prion seeding in the brains of Wisc-1 inoculated humanized mice. The heatmap indicates the percentage of positive RT-QuIC replicates out of the total of eight replicates analyzed. The scale ranges from 0 (all replicates were negative) to 1 (all replicates were positive). RT-QuIC results from WTD-Wisc-1 brain homogenates were included as a positive control, and results from age-matched non-inoculated tg650 mouse brain homogenates were included as a negative control. Mouse rPrP was used as a substrate.

RT-QuIC results are summarized in a heatmap (Figure 1B). Prion seeding activity was repeatedly detected in 77.7% of the brains of Wisc-1-inoculated mice in at least one of the six tested dilutions (Figure 1B), while 22.2% (mice #326 and #327) had inconclusive results (Figure 1B). The RT-QuIC results did not always correlate with the clinical status of the animals at the time of euthanasia. In fact, mouse #327, an animal with terminal disease (Table S1) had very poor seeding activity with only one replicate out eight being positive for two dilutions (10^-1^ and 10^-4^) while all other dilutions were strictly negative (Figure 1). In contrast, mouse #322 had strong positive seeding activity in two dilutions (10^-1^ and 10^-2^; Figure 1) yet the mouse did not develop any prion signs throughout the course of the experiment (Table S1).

Brains of all mice inoculated with the 116AG isolate were negative in RT-QuIC, regardless of the clinical state of the animals (Table S1); however, we have shown previously that this isolate induces slower disease progression, longer survival and lower seeding activity compared to Wisc-1 in cervid PrP transgenic mice ^30, 31^. None of the CWD inoculated mice, regardless of the inoculum, had seeding activity in the spinal cord and spleen.

Next, we performed western blot analysis of brain homogenates digested with proteinase K (PK; 200 µg/mL), a gold standard for the diagnosis of prion disease, to determine the presence of PK-resistant PrP^Sc^ (PrP^res^) in the brains of CWD-inoculated tg650 mice. We detected PrP^res^ in the brain of a Wisc-1-inoculated tg650 mouse, #321 (Figure 2). This mouse that was euthanized at 882 dpi upon reaching the terminal stage of disease, was the first one to show myoclonus at one year post inoculation (Table S1) and had the highest levels of seeding activity in RT-QuIC up to a dilution of 10^-6^ (Figure 1). This indicates correlation between the amount of seeding activity and PrP^res^ signal detectable in western blot. Remarkably, the PK-resistant PrP^Sc^ core presented as two fragments with estimated molecular weights of 12 – 13 and 7 – 8 kDa (Figure 2). These fragments were detected by two anti-PrP monoclonal antibodies (mAb) that recognize epitopes of the central region of PrP, Sha31 (aa 145-152) and 12F10 (aa 145-155), and by a third mAb binding an N-terminal epitope, 9A2 (aa 102-104). Notably, N-terminal mAb 12B2 (aa 93-97) only detected the 12 – 13 kDa fragment, indicating an N-terminal cleavage of this atypical PrP^Sc^ at this site (Figure 2). To rule out the possibility of a nonspecific signal, we performed a western blot with the same samples mentioned above (Figure 2) using only a secondary Ab to probe the immunoblot. The secondary Ab alone did not detect any of the PK-resistant fragments (Figure S4A), thus, validating the specificity of the PrP^res^-truncated fragments found in the brain of mouse #321 (Figure 2).

**Figure 2.**
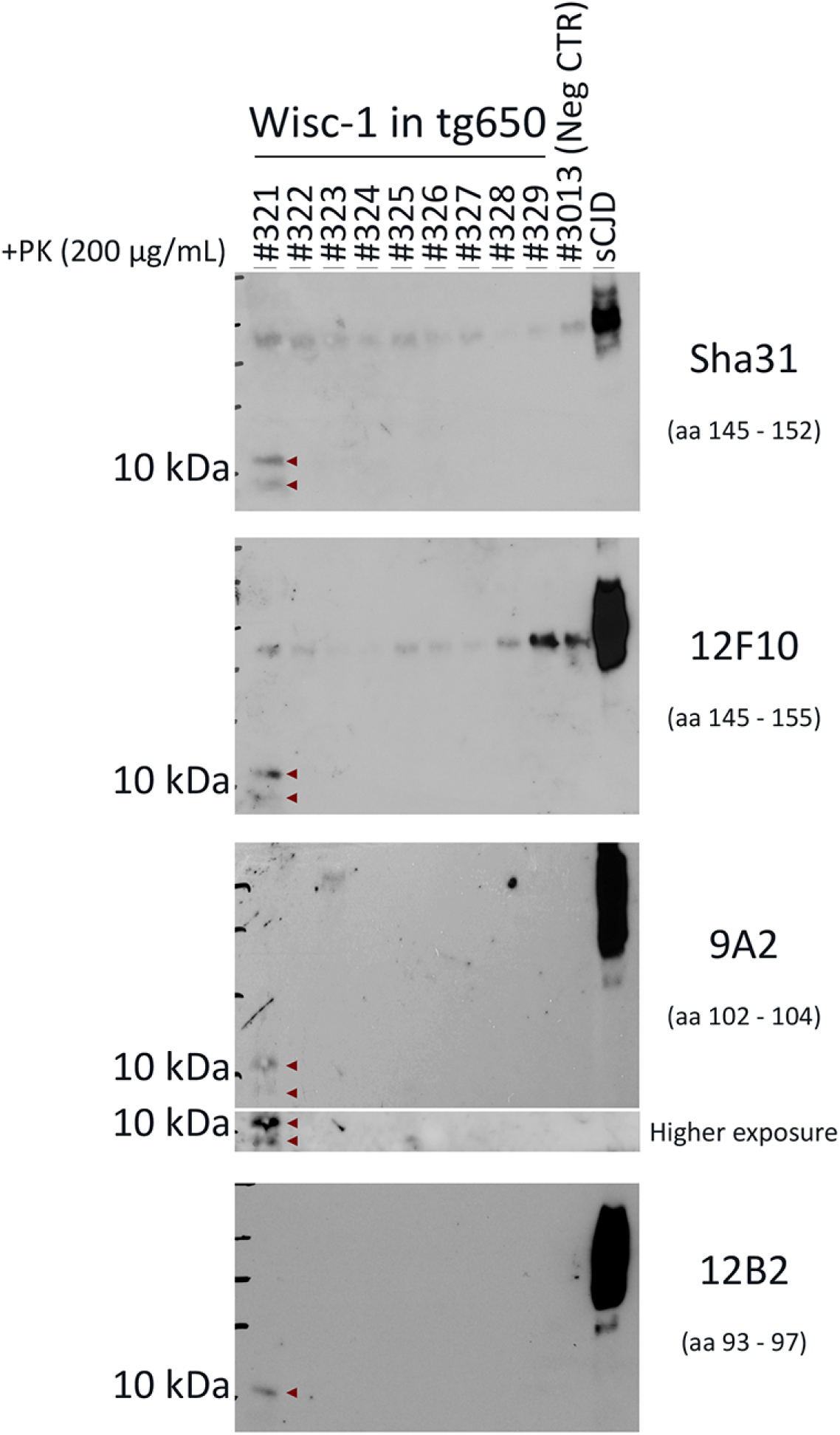
Biochemical characteristics of protease-resistant hCWD-prion protein of Wisc-1- inoculated tg650 mice. Western blot analysis of brain homogenates of tg650 mice inoculated with Wisc-1 isolate and digested with 200 µg/mL of PK using anti-PrP mAbs, from top to bottom, Sha31 (aa 145 – 152), 12F10 (aa 145 – 155), 9A2 (aa 102 – 104), and 12B2 (aa 93 – 97). A negative control, tg650 #3013, as well as a positive control (sCJD, MM1 subtype) were also included in the western blot. Higher exposure for mAb 9A2 is also shown.

Consistent with the results found in RT-QuIC, mice inoculated with 116AG isolate were all negative for PrP^res^ on western blot (Figure S5B).

We performed neuropathological analyses to assess spongiosis and abnormal PrP deposition, which are undeniable hallmarks to diagnose prion diseases. First, we analysed spongiform lesions in the gray matter of Wisc-1-inoculated tg650 mice and their age-matched negative controls. No difference in spongiform degeneration was observed between CWD inoculated animals and negative controls (data not shown). At this stage, CWD-inoculated and non-inoculated tg650 mice were old and vacuolar changes were present equally in their brains due to natural aging process. Next, we performed immunohistochemical analysis (IHC) of the brain tissues from Wisc-1- inoculated tg650 mice to detect disease-associated PrP deposits. Age-matched negative controls were used to set a baseline of normal PrP staining and determine non-disease associated PrP aggregates. Brain tissues were stained with mAb 12F10 using two different staining protocols with either PK digestion or guanidine thiocyanate denaturation (Figure 3). IHC revealed abnormal PrP deposits mainly located in the thalamus, hypothalamus, and the midbrain/pons areas of one (mouse #328) out of the six Wisc-1-inoculated mice that were tested (Figure 3). Abnormal PrP presented as granular, synaptic-like deposits surrounding the cells in these areas. Mouse #328 was euthanized at 934 dpi with subtle clinical signs (Table S1). It showed strong seeding activity in RT-QuIC (Figure 1) but was negative for PrP^res^ in western blot (Figure 2). As expected, age-matched controls were negative for abnormal PrP deposits (Figure 3 and Figure S5).

**Figure 3.**
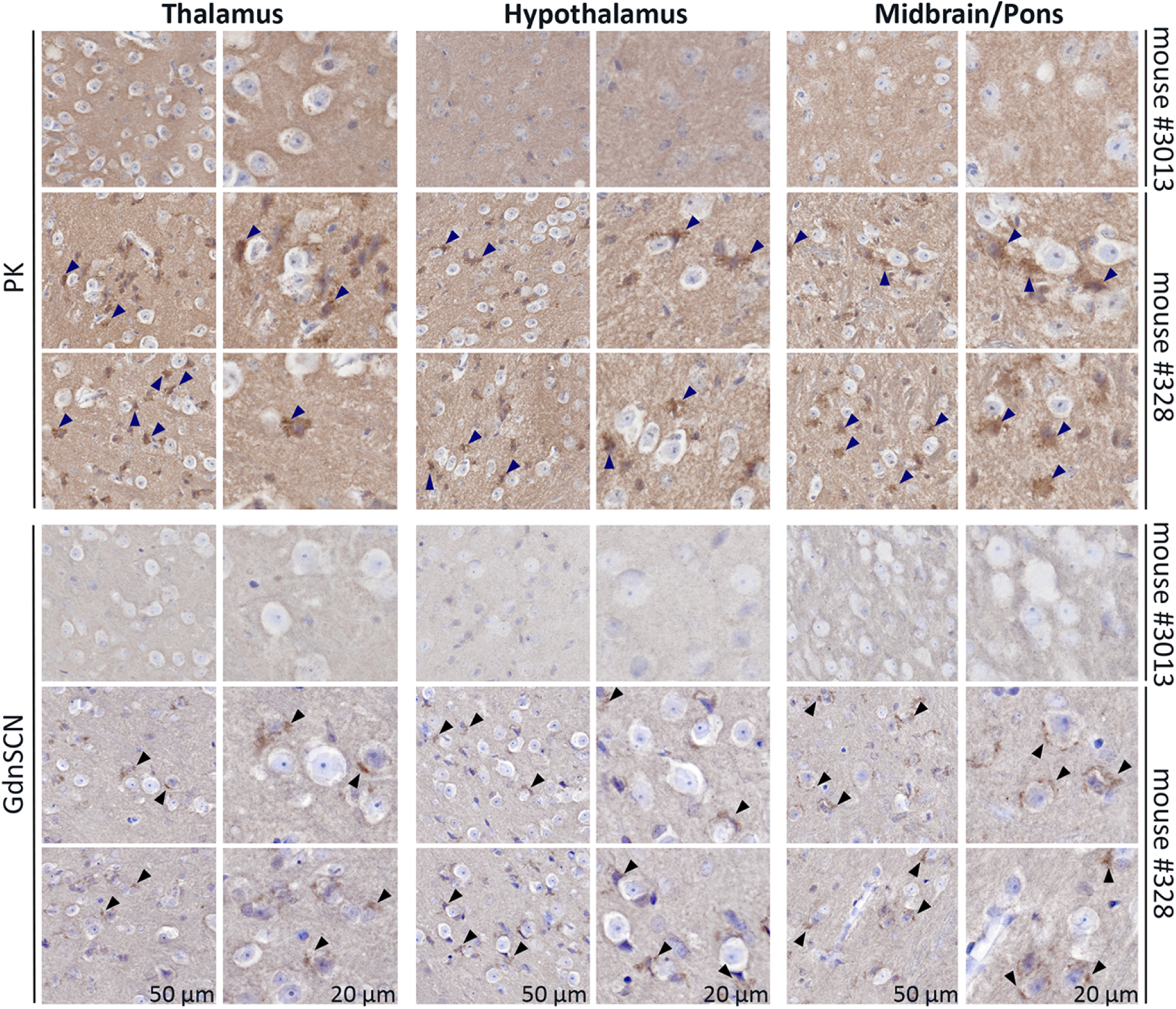
Immunohistochemistry of brain regions from Wisc-1-inoculated tg650 mice. Immunohistochemistry of Wisc-1-tg650 mouse #328, and non-infected tg650 age-matched negative control (mouse #3013) using either PK digestion (upper panels) or guanidine denaturation (GdnSCN; lower panels), using 12F10 mAb shows PrP^Sc^ deposits in the thalamus (left panels), hypothalamus (middle panels), and midbrain/pons (right panels) areas. Scale bars, 50 µm and 20 µm.

To assess infectivity and transmissibility of CWD prions in humanized mice (hCWD), we performed second passage using brain/spinal cord homogenates from mouse #327 to bank voles (n=6) and tg650 mice (n=5). The different bioassays are still ongoing, but to date, transmission of mouse #327 induced terminal prion disease at 441 dpi in one bank vole (#3054) exhibiting tremor, ataxia, kyphosis, gait abnormalities, and rigid tail. Another bank vole (#3053) died due to malocclusion at 293 dpi, yet it was monitored prior to its death because of clinical signs of transient kyphosis and gait abnormalities (dragging its left hind paw). Brain homogenates of both bank voles tested in RT-QuIC showed positive seeding activity in the brain (Figure S6A and B). However, none of the bank voles showed PrP^res^ signal in their brain homogenates in western blot.

Strikingly, second passage in tg650 mice of the brain/spinal cord homogenate pool resulted to date in terminal prion disease at 504 dpi in one mouse (#3063). This animal exhibited myoclonus at 11 months post inoculation, followed by rigid tail, rough coat, ataxia and cycles of weight loss and gain. Western blot analysis of PK-digested (200 µg/mL) brain homogenates revealed atypical PrP^res^ fragments (Figure 4), resembling those of Wisc-1-inoculated tg650 mouse #321 (Figure 2) in molecular weights and N-terminal cleavage site. These results demonstrate the presence of transmissible prion infectivity (hCWD) in the CNS of terminally sick tg650 mouse #327, despite the absence of detectable seeding activity in RT-QuIC (Figure 1), PrP^res^ in western blot (Figure 2), and abnormal PrP deposits in IHC (Figure S5).

**Figure 4.**
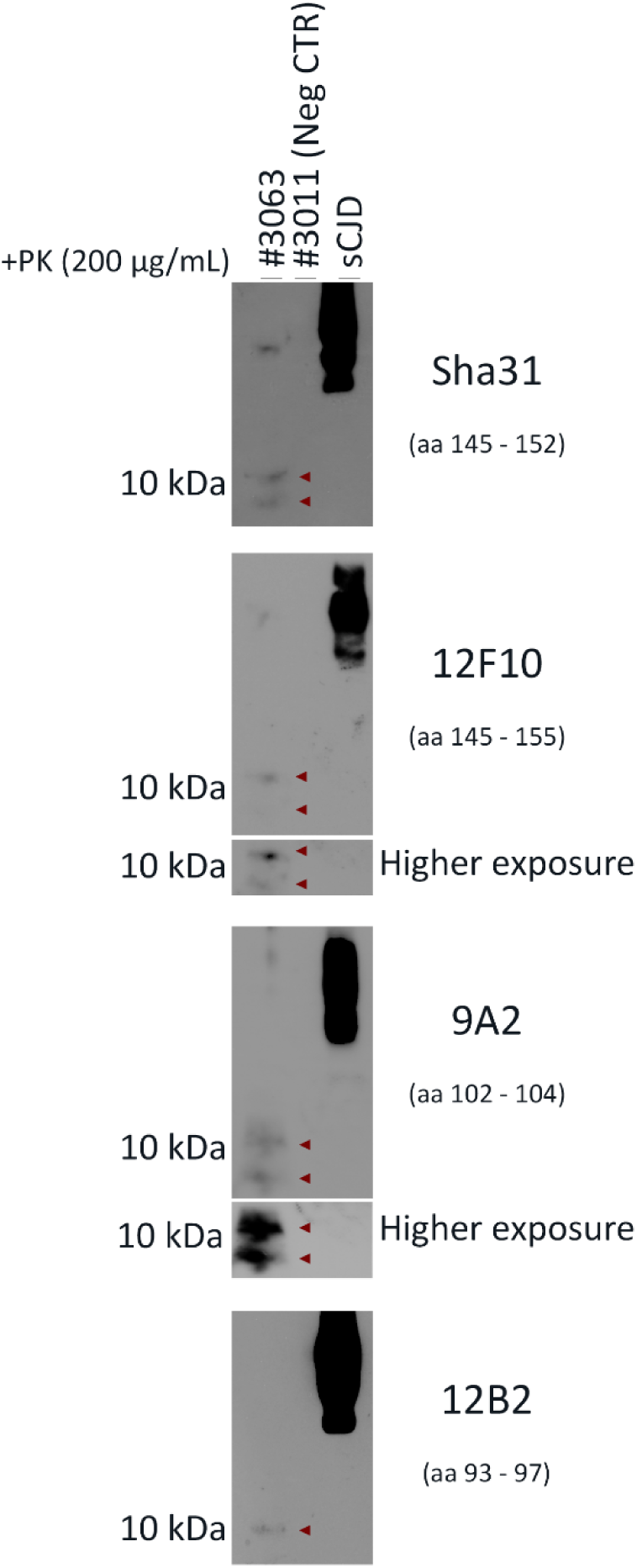
Transmission of CWD-tg650 brain/spinal cord homogenates from mouse #327 to tg650 mice. Western blot analysis of brain homogenates from tg650 mouse (#3063; 2^nd^ passage) inoculated with brain/spinal cord homogenates from tg650-Wisc-1 #327 (1^st^ passage) and digested with 200 µg/mL of PK using anti-PrP mAbs, from top to bottom, Sha31 (aa 145 – 152), 12F10 (aa 145 – 155), 9A2 (aa 102 – 104), and 12B2 (aa 93 – 97). A negative control, tg650 #3011, as well as a positive control (sCJD) were included in the western blot. Higher exposures for mAbs, 12F10 and 9A2, are also shown.

### CWD inoculated humanized mice shed infectious prions in feces

CWD prions are excreted in feces of infected, preclinical cervids ^5,35, 36^. Therefore, we collected feces from symptomatic and asymptomatic Wisc-1 or 116AG-inoculated tg650 mice, as well as their age-matched controls, between 600 dpi – 750 dpi. We analysed fecal homogenates by RT-QuIC. We used mouse recombinant PrP (rPrP) as a substrate and, as for the brain homogenates, substrate replacement for enhanced sensitivity ^35, 37^. To verify whether fecal contents may contain inhibitors of amyloid formation we performed spiking experiments. We used fecal homogenates from age-matched non inoculated controls to spike with sCJD brain homogenates as additional positive control (Figure S3B). Remarkably, 50 % of the mice showed positive seeding activity in some replicates and dilutions (Figure S7A). Wisc-1-inoculated mouse #327 had consistently high levels of seeding activity with 75% – 100% positive replicates, up to a dilution of 10^-3^ (Figure 5). We confirmed these results using human rPrP (Figure S7B).

**Figure 5.**
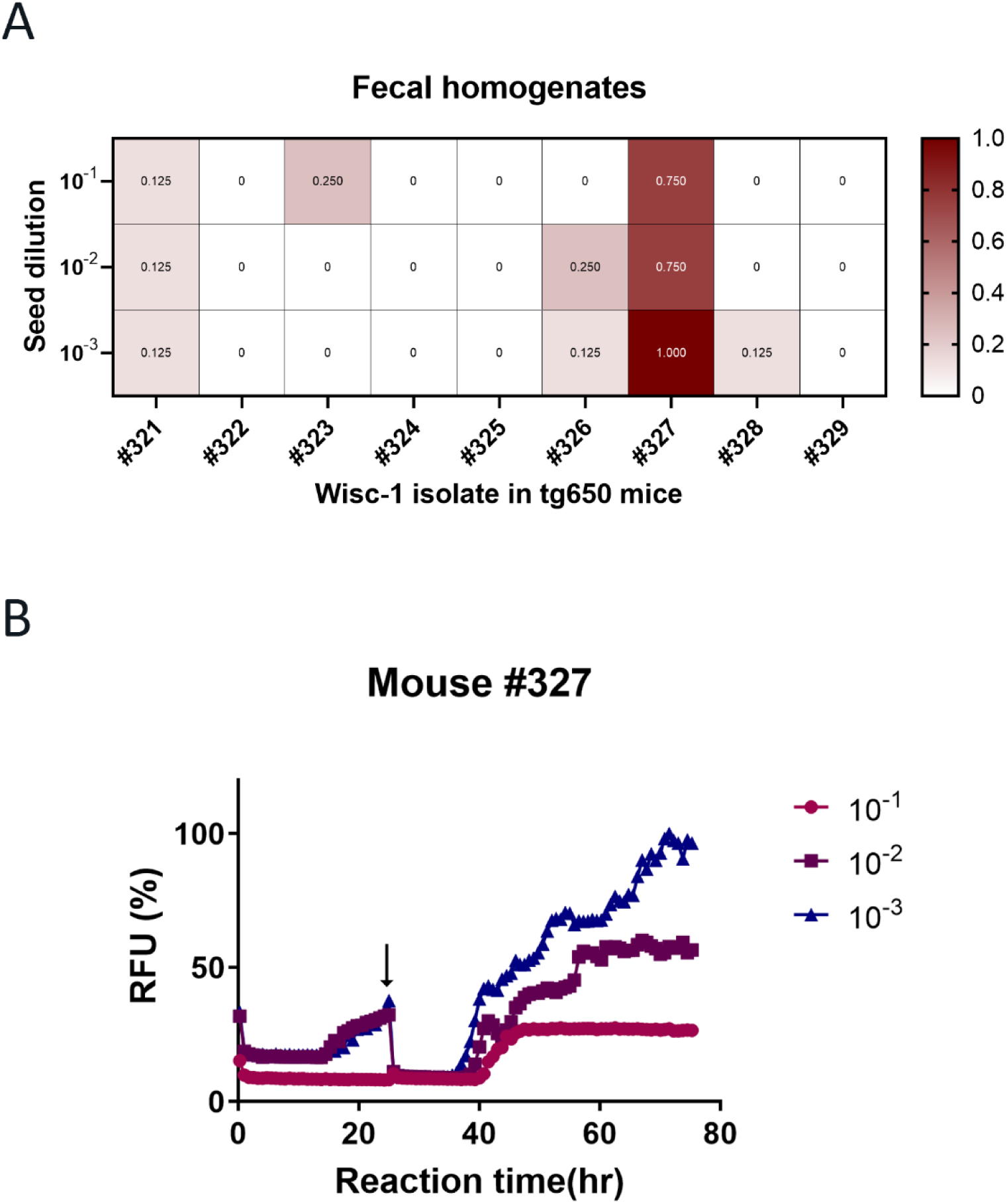
Prion seeding activity in feces of Wisc-1-inoculated humanized mice. A. Summary of RT-QuIC analysis of prion seeding in fecal homogenates of Wisc-1 inoculated tg650 mice. **A.** The heatmap indicates the percentage of positive RT-QuIC replicates out of the total of eight replicates analyzed. The scale goes from 0 (all replicates were negative) to 1 (all replicates were positive). **B.** The graphs depict a representative RT-QuIC assay of fecal homogenates from tg650 mouse #327. Fecal homogenates were serially diluted (10^-1^ to 10^-3^). Mouse rPrP was used as a substrate. Replacement with fresh assay buffer containing rPrP is indicated with a black arrow.

To validate the presence of infectious prions in fecal homogenates, we intracerebrally inoculated bank voles (n=9) and tg650 mice (n=10) with 10% sonicated fecal homogenates from mouse #327 (summarized in Figure S1). Fecal homogenate inoculation to tg650 mice caused, so far, prion disease in 50% of the inoculated mice with the bioassay still ongoing. The mice presented with myoclonus at an early stage of the course of the experiment, followed by cycles of weight loss and gain, and finally ataxia. These mice were euthanized upon reaching terminal stage of the disease, between 534 – 630 dpi. However, analyses of brain homogenates from these mice in western blot were negative for PK-resistant PrP^Sc^. To date, fecal homogenate inoculation resulted in prion disease in bank voles (Figure 6A), with varying survival times (198 dpi to 531 dpi) and with an attack rate of 66.6% (6/9 voles). Most of the euthanized voles presented subtle clinical signs (5/6), with ataxia as a predominant symptom, and were euthanized because of excessive weight loss. One bank vole (#3430) presented with progressive weight loss, kyphosis, ataxia, gait abnormalities and was euthanized upon reaching terminal disease stage at 241 dpi. In all tested brain homogenates from bank voles inoculated with Wisc-1-tg650 fecal homogenates, seeding activity was detectable by RT-QuIC (Figure S6C-E). In addition to the high levels of seeding activity found in the brain homogenates of bank vole #3430 (Figure S6E), spinal cord homogenates from this animal also showed strong seeding activity (Figure S6F).

**Figure 6.**
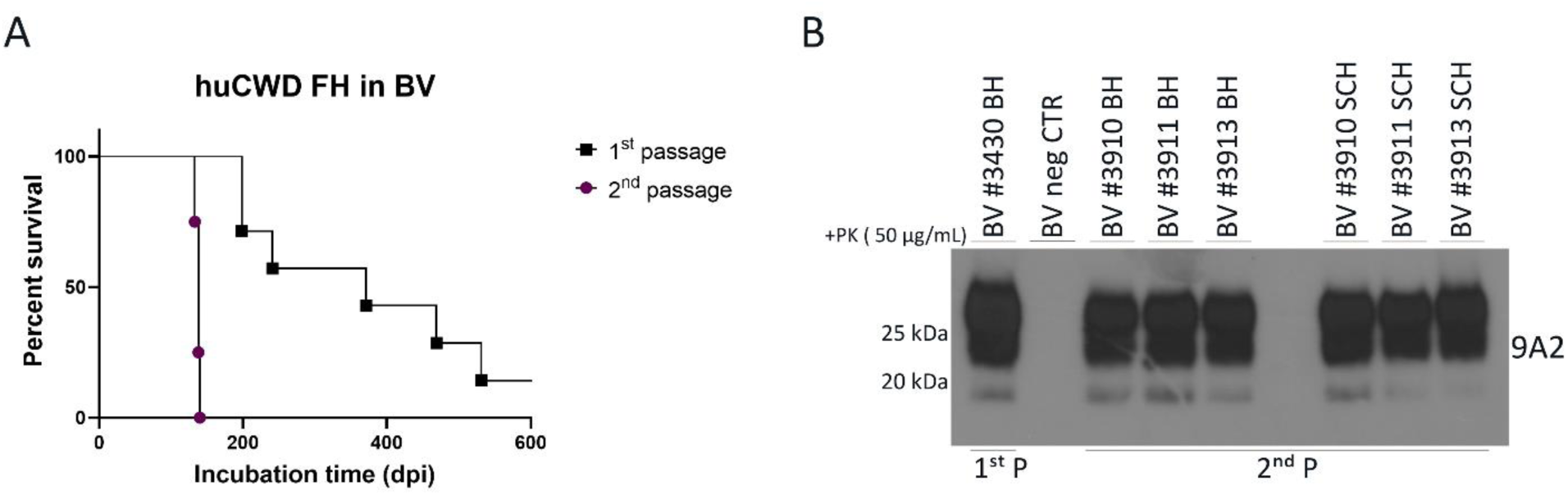
Transmission of CWD-tg650 fecal material to bank voles. 10% fecal homogenates from Wisc-1-inoculated tg650 mouse #327 was inoculated intracerebrally to bank voles (1^st^ passage), and brain homogenates of 1^st^ passage bank was transmitted to the same host (2^nd^ passage). **A.** A Kaplan-Meier curve depicting the survival times of bank voles inoculated with Wisc-1-tg650 fecal homogenates upon 1^st^ and 2^nd^ passage. **B.** Western blot analyses of bank voles inoculated with Wisc-1-tg650 fecal homogenates, 1^st^ passage (lane 1, brain homogenates), and 2^nd^ passage (brain homogenates, lanes 3-5, and spinal cord homogenates, lanes 7-9). Samples, as well as a bank vole brain homogenates as a negative control, were digested with 50 µg/mL of PK. The western blot was probed with mAb 9A2.

Next, we performed western blot analysis of PK-digested (50 µg/mL) brain and spinal cord homogenates from bank vole #3430. We detected a typical three banding pattern-PrP^res^ signal in both tissues (Figure 6B and Figure S8A).

Remarkably, analyses of small intestine and colon homogenates of bank vole #3430 by RT-QuIC revealed positive seeding activity (Figure S6G and H, respectively). Small intestine homogenates displayed stronger seeding activity up to a dilution of 10^-5^ compared to colon homogenate, possibly due to the presence of inhibitory components ^35^ in the colon or generally lower levels of prion seeding activity. However, after PK digestion, none of these tissues was positive for PrP^res^ in western blot. Spleen homogenates of fecal homogenate-inoculated bank voles were negative in both western blot and RT-QuIC.

Next, to validate the transmissibility of prions present in bank voles inoculated with Wisc-1-tg650 fecal homogenates, we performed a second passage of brain homogenates from bank vole #3430 into the same host (n=4). Second passage resulted in a complete attack rate (Figure 6A), and all inoculated bank voles succumbed to terminal prion disease. The decrease in survival time upon second passage (average of 137±3 dpi; Figure 6A) illustrates an adaptation of these prions to their new host. Western blot analyses of samples from second passage showed the presence of a three banding PrP^res^ pattern in the brain and spinal cord homogenates of these animals (Figure 6B). As a negative control, we also inoculated bank voles with fecal homogenates of tg650 non-inoculated mice. One of the voles was euthanized at 550 dpi to test for seeding activity by RT-QuIC, and it was found negative. To date (>600 dpi), the remaining bank voles in this control group are still healthy and not displaying any signs.

Next, we wanted to compare PK-resistant PrP^Sc^ biochemical signature in brain homogenates of bank voles inoculated with Wisc-1-tg650 fecal homogenates, Wisc-1 original deer isolate, and Wisc-1 inoculated in bank voles ^31^ (bvWisc-1; 1^st^ and 2^nd^ passage). Western blot analyses showed that PrP^res^ signal in brain homogenates of bank voles inoculated with Wisc-1-tg650 fecal homogenates (Figure 6B, 7A and Figure S8A) differed from PrP^res^ signal in brain homogenates of bvWisc-1 (Figure 7A and Figure S8A). The unglycosylated PrP^res^ band of bvWisc-1 migrates at an approximate molecular weight of 20 kDa, different from that of the Wisc-1 original deer isolate (18 – 19 kDa; Figure 7A and ^31^). Interestingly, unglycosylated PrP^res^ in brain and spinal cord homogenates from bank vole #3430 inoculated with Wisc-1-tg650 fecal homogenates migrated at 18 – 19 kDa (Figure 6B, 7A and Figure S8A) resembling that of the Wisc-1 original deer isolate (Figure 7A). When we quantified the signal of the di-, mono-, and unglycosylated bands (Figure 7B and Figure S8B), we found that bvWisc-1 presented with a predominant di-glycosylated band and a non-detectable unglycosylated band, compared to Wisc-1 deer isolate and bank voles inoculated with Wisc-1-tg650 fecal homogenates in 1^st^ and 2^nd^ passage (Figure 7B and Figure S8B). The PrP^res^ signal intensity in bank vole #3430 was stronger compared to that of bvWisc-1, considering that PK digestion was performed at different concentrations, 25 µg/mL for bvWisc-1 compared to 50 µg/mL for bank vole #3430, and different volumes were applied for western blot analysis, with 20 times higher volume loaded for bvWisc-1 compared to bank vole #3430 (Figure 7A and Figure S8A). These data demonstrate that humanized tg650 mice inoculated with CWD prions shed prion infectivity into feces that are able to generate transmissible PrP^Sc^ in bank voles.

**Figure 7.**
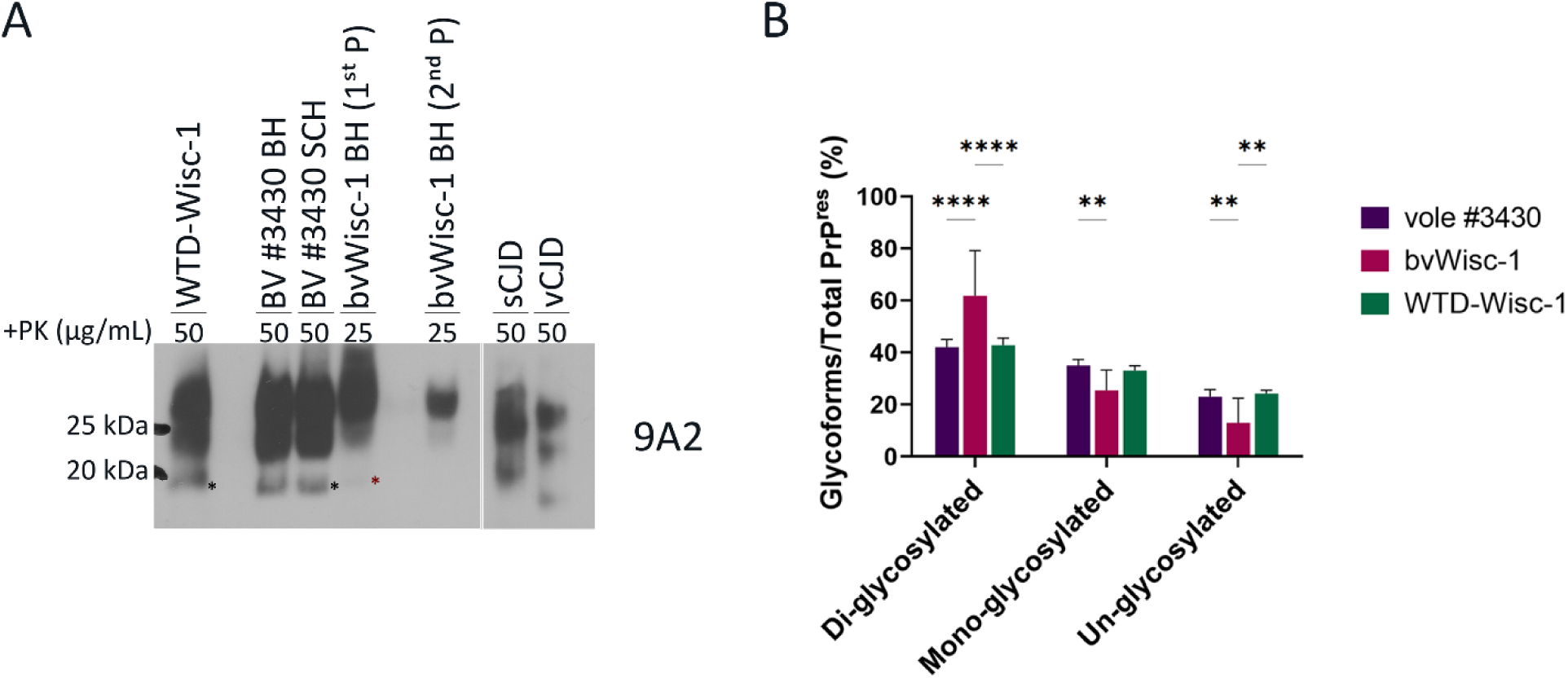
Biochemical characteristics of protease-resistant PrP^Sc^ in bank voles inoculated with fecal homogenates from Wisc-1-inoculated tg650 mice. **A.** Western blot analyses of bank voles inoculated with fecal homogenates from Wisc-1-inoculated tg650 (lane 3 and 4, brain and spinal cord homogenates, respectively) and Wisc-1 deer isolate (lane 1) digested with 50 µg/mL of PK, as well as Wisc-1 passaged in bank voles (bvWisc-1), 1^st^ (lane 5) and 2^nd^ (lane 7) passage digested with 25 µg/mL of PK. sCJD and vCJD were loaded as positive controls and were digested with 50 µg/mL of PK. The western blot was probed with mAb 9A2. The amount of bvWisc-1 loaded on the gel was 20x that of bank vole #3430. **B.** Quantification of the glycoform ratio of Wisc-1 isolate, fecal homogenate inoculated bank vole (1^st^ and 2^nd^ passage), and bvWisc-1 (1^st^ and 2^nd^ passage) prions. Statistical analyses were performed using two-way ANOVA (Tukey’s multiple comparisons test).

## Discussion

Our findings strongly suggest that CWD is an actual public health risk. Here, we use humanized mice to show that CWD prions can cross the species barrier to humans, and remarkably, infectious prions can be excreted in feces.

Many factors have to be taken into account for prions to be able to cross the species barrier. Notably, the homology between the host PrP primary structure and the donor PrP^Sc^ plays a crucial role in species barrier ^38–41^. In addition, the existence of different CWD strains and the impact of the *Prnp* polymorphism, the extensive incubation period, and the atypical clinical presentation should be considered when investigating the risk of CWD. Determining that CWD poses a significant risk to human health is of a crucial importance to prevent a major public health and economic crisis similar - if not greater - to that of BSE transmission to humans (i.e., vCJD).

Several studies have evaluated the susceptibility of humans to CWD ^42^. Most *in vitro* studies had found that CWD prions are capable to convert human PrP^C^, however, either the efficiency of the conversion was low or relied on protein modifications by denaturation ^17, 18, 21, 43, 44^. RT-QuIC assay showed that PrP^CWD^ can seed the conversion of human recombinant PrP, however, PrP^BSE^ did not seed it making the veracity of these results questionable ^45^. Transgenic mouse models overexpressing the human prion protein have been extensively used to assess the transmission of various human prions *in vivo*, thus, they are also used as a model to test the zoonotic potential of prions from other species such as cervids (CWD) ^10–12, 14, 15, 19, 20, 46^. Different transgenic lines with different polymorphism at residue 129 (M or V), expressing human PrP^C^ at different expression levels, and in variable tissues (because of the transgene construct), were used in these studies, that were infected with different sources of CWD prions.

CWD transmission to humanized mice was assessed in all these studies by monitoring the clinical signs, examining the brain for neuropathology associated with prion disease, and IHC staining for PrP deposits. Brains were also analyzed by western blot after PK digestion for detection of PrP^res^. Wilson and coworkers used an antigen immunoassay, to screen for PrP^CWD^ ^19^, and Race *et al.* used RT-QuIC to assess the PrP amyloid formation in the brain of CWD inoculated humanized mice ^15^. In three of the *in vivo* studies ^11, 15, 19^, clinical signs were observed in a relatively small number of CWD inoculated mice. Race *et al.* showed positive seeding activity in the brains of very few suspect animals, however, these results were inconsistent, and the authors questioned their significance ^15^. Prion disease could not be detected in any of the other studies. As RT-QuIC assay was not available at the time of other reports ^4,10–12^, they might have missed subclinical manifestations in their CWD-inoculated humanized mice. In addition, except for one study where the mice were kept up to 798 dpi ^15^, in most of the studies the experiments were terminated between 500 – 600 dpi. In our study, tg650 mice were kept for up to >900 dpi, which is at least +15% of the survival compared to the longest study. This could be a significant asset to this line when it comes to human prion diseases with a very prolonged incubation period.

In our study, positive seeding activity observed in the brain of 77.7% of Wisc-1-tg650 inoculated mice was consistent and reproducible, however, one concern was that these results are the product of remaining inoculum as reported by Martin *et al.* ^47^, or a combination of both *de novo* generated PrP^CWD^ in these mice, with regards to our results, and remnant inoculum. Nevertheless, our results are in disfavor of a scenario involving a remnant inoculum alone. In fact, transmission of brain homogenates from knockout mice with remaining sCJD inoculum generated prion disease in tg650 mice that was undistinguishable from that generated by sCJD inoculation of these mice. In our case, transmission of Wisc-1-CWD to tg650 mice resulted in clinical disease, and transmission of hCWD occult prions (mouse #327) that were transmissible upon second passage exposing an atypical PrP^res^ pattern (mouse #3063) similar to that detected in first passage (mouse #321). The presence of infectious prions shed in feces is an argument in favor of the existence of *de novo* generated PrP^Sc^ in these mice. Another argument is the fact that bank voles inoculated with material from Wisc1-inoculated tg650 had a different disease manifestation compared to bank voles inoculated with Wisc-1 deer isolate (Figure 7 and Figure S8). This implicates that prions generated following Wisc-1 inoculation in tg650 mice underwent a modification governed by the presence of the human *PRNP* sequence. The recent claim by Wang *et al.* of susceptibility of humanized mice to CWD failed to transmit the original elk isolate to the mice, rather, they generated PrP^CWD^ by PMCA using V129 PrP^C^ as substrate ^20^. The study showed that CWD prions only converted V129-PrP^C^ but not M129-PrP^C^, somehow contradicting previous reports demonstrating that CWD also converted M129-PrP^C^ ^43^.

Our results indicate that if CWD crosses the species-barrier to humans, it is unlikely to resemble the most common forms of human prion diseases with respect to clinical signs, tissue tropism and PrP^Sc^ signature. For instance, PrP^Sc^ in variable protease sensitive prionopathy (VPSPr), a sporadic form of human prion disease, and the genetic form Gerstmann-Sträussler-Scheinker syndrome (GSS) is defined by an atypical PK-resistant PrP^Sc^ fragment that is non-glycosylated and truncated at both C and N termini, with a molecular weight between 6 and 8 kDa ^48–51^. These biochemical features are unique and distinctive from PrP^Sc^ (PrP27-30) found in most other human or animal prion disease. The atypical PrP^Sc^ signature detected in brain homogenate of tg650 mice #321 (1^st^ passage) and #3063 (2^nd^ passage), and the 7 – 8 kDa fragment (Figure 2 and 4) are very similar to that of GSS, both in terms of migration profile and the N-terminal cleavage site.

CWD in humans might remain subclinical but with PrP^Sc^ deposits in the brain (e.g., mouse #328; Figure 3), clinical with untraceable abnormal PrP (e.g., mouse #327) but still transmissible and uncovered upon subsequent passage (e.g., mouse #3063), or prions have other reservoirs than the usual ones, hence the presence of infectivity in feces (e.g., mouse #327) suggesting a potential for human-to-human transmission and a real iatrogenic risk that might be unrecognizable. Here, humanized mice inoculated with CWD deer isolates had an atypical onset of the disease with myoclonus (93.75%), before presenting typical clinical signs, generating prions that presented with either atypical biochemical signature (#321 and #3063), shed in feces (#327), or were undetectable by the classical detection methods. The fact that we could not establish a strong correlation between disease manifestation in tg650 mice inoculated with Wisc-1- or 116AG-CWD and the presence of abnormal PrP (western blot, IHC or RT-QuIC) might be explained by the presence of heterogeneous protease sensitive and resistant PrP^Sc^ in the brains of infected mice with different seeding properties *in vitro*. Indeed, such heterogeneity and distinct seeding activities and infectivity of abnormal PrP fragments was observed in VPSPr cases ^52, 53^.

The finding that infectious PrP^Sc^ was shed in fecal material of CWD-infected humanized mice and induced clinical disease, different tropism, and typical three banding pattern-PrP^res^ in bank voles that is transmissible upon second passage is highly concerning for public health. The fact that this biochemical signature in bank voles resembles that of Wisc-1 original deer isolate and is different from that of bvWisc-1, in the migration profile and the glycoform-ratio, is valid evidence that these results are not a product of contamination in our study. If CWD in humans is found to be contagious and transmissible among humans, as it is in cervids ^5^, the spread of the disease within humans might become endemic. In contrast to bank voles inoculated with fecal homogenates from mouse #327, so far, we could not detect a resistant-PrP^Sc^ fragment in the brain homogenates of fecal homogenates inoculated-tg650 mice. The presence of PrP^res^ in these mice will allow us to determine if the molecular signature of hCWD prions from the brain (mouse #321 and #3063) vs feces are the same. Previously, Beringue *et al.* found that extraneural prions, compared to neural prions, helped greater to overcome the species barrier to foreign prions, in addition, different strain types emerged from such serial transmission ^54^. Our data also suggest that prions found in the periphery may hold higher zoonotic potential than prions found in neural tissues. In fact, upon second passage, 50% of the tg650 mice inoculated with fecal homogenates from mouse #327 had succumbed with terminal disease compared to only 20% of brain/spinal cord homogenates inoculated-tg650 mice suggesting that hCWD prions found in feces transmit disease more efficiently. Our results also suggest that epidemiological studies ^4^ may have missed subclinical and atypical infections that are/might be transmissible, undetected by the gold standard tests, i.e., western blot, ELISA, and IHC.

Ideally, an oral inoculation of mice with CWD prions would have mimicked best the natural route of exposure to acquired prion diseases like CWD. However, in our case, an oral inoculation would have been impossible considering the lifespan of the rodent models. It took us an extended amount of time, over two years and a half, to have a better idea of the extent of CWD-transmission in the tg650 model, even with intracerebral inoculation, usually the fastest way to induce prion disease in a rodent model. Taking this into consideration, our study is the strongest proof-of-principal that CWD is transmissible to humans. Using humanized mice, we demonstrated the zoonotic potential of CWD. Furthermore, our findings provide striking insights into how CWD might manifest in humans and the impact it may have on human health. We have used Wisc-1/CWD1, one of the most common CWD strains, notably WTD prions, which have been shown to be more prone to generate human prions *in vitro* ^43^. This implies a high risk of exposure to this strain, e.g., through consumption or handling of infected carcasses, in contrast to rarer CWD strains, and therefore, an actual risk for human health. In addition, CWD surveillance in humans should encompass a wider spectrum of tissues/organs tested and include new criteria in the diagnosis of potential patients.

## Methods

### Ethics statement

This study had the approval of the Canadian Human Ethics Board for the use of CJD isolates (protocol number REB18-0505), and strictly followed the guidelines of the Canadian Council for Animal Care. All experiments detailed in the study were performed in compliance with the University of Calgary Animal Care Committee under protocol number AC18-0047. Prior to inoculation and euthanasia, isoflurane was used as anesthetic at a concentration of 5% (flow rate of 0.8 L/min) for induction, and then lowered to 0.5–1% for maintenance of general anesthesia during the procedure.

### Prion material

Prion isolates were prepared separately as 20% (w/v) brain homogenates in phosphate-buffered saline pH 7.4 (PBS; Life Technologies, Gibco) using the MP Biomedicals fast prep-24 homogenizer (Fisher). Aliquots were stored at −80°C until further use. Homogenization and aliquoting of the CWD isolates was done in two different laboratories, and during handling, gloves were changed between CWD isolates to avoid any cross contamination. Wisc-1 deer isolate was obtained upon experimental infection of WTD of wild-type *Prnp* genotype (QQ95, GG96, and AA116) orally dosed with CWD inoculum from hunter harvested deer ^55^. This isolate was characterized as Wisc-1 strain ^32, 33, 56^. WTD-116AG isolate was a field isolate from an animal with a polymorphic *Prnp* genotype (QQ95, GG96, and AG116) provided by the Canadian Wildlife Health Cooperative (CWHC), Saskatoon, SK, Canada ^30^. It was from a 5 year old wild male WTD that was reported to exhibit clinical signs (wasting syndrome) and was tested positive for CWD after its death in the field. Except for residue 116, no other polymorphisms were found in the *Prnp* gene of this animal ^30^. As a positive control for western blots, we used MM1-sCJD and MM2b-vCJD brain tissues kindly provided by Dr. Stéphane Haïk (Paris Brain Institute, France) that were handled in a separate laboratory.

Fecal homogenates were prepared according to a well-established protocol in our lab ^35^. Briefly, fecal pellets collected from CWD-inoculated tg650 mice (between 600 dpi – 750 dpi), and age-matched controls were weighed and prepared in fecal extraction buffer composed of 20 mM sodium phosphate (pH 7.1), 130 mM NaCl, 0.05% Tween 20, 1mM PMSF and 1X complete protease inhibitors (Roche) at a final concentration of 20% (w/v). Fecal pellets were homogenized in the MP Biomedicals fast prep-24 homogenizer and placed on a rotary shaker for 1 hour at room temperature, then centrifuged at 18,000 x g for 5 minutes, supernatants were collected. Fecal homogenate (1 mL) was mixed with N-lauryl-sarcosine at a final concentration of 2% and incubated at 37° C with constant shaking at 1,400 rpm for 30 minutes. The samples were adjusted to 0.3% sodium phosphotungstic acid (NaPTA) by adding a stock solution containing 4% NaPTA and 170 mM magnesium chloride (MgCl_2_), pH 7.4, incubated for 2 hours at 37° C with constant shaking and centrifuged for 30 minutes at maximum speed (15,800 x g) at room temperature. Pellets were washed using cell lysis buffer containing 0.1% N-lauryl-sarcosine, and then resuspended in 1/10 of the original sample volume in RT-QuIC dilution buffer.

### Animal study

All animal transmissions in tg650 and bank vole models are summarized in a scheme (Figure S1). We used tg650 mice, a transgenic mouse line that overexpresses human PrP^C^ (MM129) approximately 6 fold ^29^. The lifespan of these animals is about 2.5 years, and they are not known to develop any spontaneous prion disease ^29, 57^. We intracerebrally (i.c.) inoculated ten 6 – 8-week-old tg650 females with 1% brain homogenates from CWD-positive WTD (Wisc-1 and 116AG), or a pool of 1% brain/spinal cord homogenates from first passage tg650-Wisc-1 (mouse #327) into the right parietal lobe using a 25-gauge disposable hypodermic needle. Alongside the CWD-inoculated tg650 mice, we also had ten tg650 age-matched controls. Strict guidelines were followed to avoid cross contamination between the CWD isolates during the inoculation procedure and throughout the course of the experiment. No human prion infected materials were handled in the animal facility where the inoculated tg650 mice were housed. We also inoculated bank voles expressing a methionine at position 109 of the *Prnp* gene ^58, 59^. Two to three months old male and female bank voles were i.c. inoculated with a pool of 1% brain/spinal cord homogenates (n= 6), or 10% fecal homogenates (n=9) that were sonicated in 2-seconds burst of sonication (∼130 - 170 watts; QSonica Q700 sonicator) followed by a rest time of 1-second for 8 minutes. Sonication was used to damage nucleic acids and inactivate bacteria and viruses with minimal effects on prion titers. For second passage a set of 4 bank voles was inoculated with brain homogenates from bv #3430. Inoculated animals were initially monitored once a week. Upon onset of clinical signs, in this particular and atypical case, myoclonus, they were monitored daily. At terminal stage of disease, clinical mice and bank voles were exhibiting rigid tail, rough coat, gait abnormalities, ataxia, kyphosis, and cycles of weight loss and gain. We defined animals with subtle clinical signs as those that developed some subtle and/or transient signs that did not progress during the course of the experiment; however, all of these animals had the myoclonus. Animals with terminal clinical signs were those with confirmatory prion signs that did progress and reached terminal stage of disease. At experimental endpoints, animals were anaesthetized and then euthanized by CO_2_ overdose. After perfusion of animals, brains were collected and either fixed in formalin or frozen at -80°C. Experimental termination endpoints were pre-determined for tg650 mice (>900 dpi). One mouse was found dead at 213 dpi and, therefore, was excluded from the analyses of Wisc-1 inoculated tg650 mice (#330). We also excluded three animals from the analyses of 116AG-inoculated tg650 mice, two that were found dead (#335 and #337) and one which was euthanized because of a humane endpoint (#334).

### Neuropathology and immunohistochemistry (IHC)

Fixed sagittal sections of brains from six CWD-inoculated animals and two of their age-matched counterparts (negative controls) were paraffin embedded. Serial sections of 5 μm thickness were cut and stained using hematoxylin and eosin (H&E; Leica) to evaluate spongiform changes. For IHC, for a first set of slides we used guanidine thiocyanate (GdnSCN) denaturation. Sagittal sections were pre-treated with high-pressure autoclaving (2.1 × 105 Pa) for 30 minutes in citric acid (10 mM), pH 6.0, at 121° C, followed by treatment with 98% formic acid for 10 minutes and 4 M GdnSCN for 2 h at room temperature. For a second set of slides, we used PK digestion instead of GdnSCN denaturation. The slides were treated with 89% formic acid for 10 minutes, digested using 4 μg/mL of PK for 5 minutes at 37° C, followed by pre-treatment with high-pressure autoclaving (2.1 × 105 Pa) for 30 minutes in citric acid (10 mM), pH 6.0, at 121°C. Abnormal PrP accumulation was examined using a commercially available ARK (Animal Research Kit)/HRP kit (DAKO) by using the anti-PrP monoclonal antibody 12F10 (1:100; Cayman) for 30 minutes at 37° C and sections were counterstained with hematoxylin. Slides were scanned using the Olympus VS110-5S scanner and images were analyzed using OlyVIA software (Olympus). All images were treated in a similar manner.

### PrP^res^ western blot detection

For PrP analysis in tg650 brains, brain homogenates (20%) prepared in PBS from different animals, including negative controls, were mixed with an equal volume of 100 mmol/L Tris-HCl (pH 7.4) – 2% sarkosyl for 30 minutes prior to PK (200 μg/mL; Roche) digestion for 1 h at 37° C. The reactions were terminated by adding 1X pefabloc proteinase inhibitor (Roche). Samples were boiled in sample buffer at 100° C for 10 minutes. For bank vole brain extracts, 20% brain homogenates in 100 mmol/L Tris-HCl (pH 7.4) – 2% sarkosyl were incubated for 1 h at 37° C with PK at a final concentration of 25 or 50 µg/mL. Protease treatment was stopped by adding 1X pefabloc and an equal volume of isopropanol/butanol (1:1 vol/vol) was added to the samples, which were then centrifuged at 20,000 × *g* for 10 minutes. The pellets were resuspended in denaturing sample buffer and heated for 10 minutes at 100° C. Fifty μg of protein were loaded, unless otherwise stated and separated using precast 12% Bis-Tris gels (ThermoFisher), and then electrophoretically transferred to PVDF membranes (GE Healthcare Life). PVDF membranes were blocked for 1 h in PBS-Tween (0.1%) containing skim milk powder (5%) and probed using anti-PrP monoclonal antibodies Sha31 (1:15000; Cayman), 12F10 (1:200; Cayman), 9A2, and 12B2 (1:1000; Wageningen Bioveterinary Research) overnight at 4° C. Washing steps were performed using PBS-Tween buffer followed by horseradish peroxidase-conjugated goat anti-mouse IgG antibody (Sigma-Aldrich) for 30 minutes at room temperature. A separate blot was probed using only horseradish peroxidase-conjugated goat anti-mouse IgG antibody for 30 minutes at room temperature to assess the specificity of the bands detected with primary mAbs (Supplementary Fig. 3). Signals were developed using ECL-plus detection (Millipore). Images were acquired on X-ray films (Denville Scientific). For calculation of the glyco-form ratios, Image J software was used to quantify and determine the relative values of PrP^res^ signals (n≥ 4).

### Preparation of recombinant PrP (rPrP) substrate

The mature form of mouse, human or bank vole (aa23-231) PrP was cloned into pET41a expression vectors (EMD Biosciences) and expressed in *E. coli* Rosetta using the Express Autoinduction System (Novagen). Inclusion bodies were prepared using the Bug Buster reagent (Novagen) and solubilized in lysis buffer (guanidine-HCl 8 M (Sigma-Aldrich)), sodium-phosphate 100 mM, Tris-HCl 10 mM, pH 8.0 (Sigma-Aldrich)) for 50 minutes at 23° C and then centrifuged at 16 000 g for 5 minutes at 23° C. Binding, refolding, and elution using an AKTA Explorer system has been described previously ^60^.

### RT-QuIC assay

For brain, spinal cord, small intestine, colon, and fecal homogenates, real-time QuIC was performed as described previously ^30, 35^. Briefly, reactions were set up in assay buffer containing 20 mM sodium phosphate (pH 6.9; Sigma-Aldrich), 300 mM NaCl (Sigma-Aldrich), 1 mM EDTA (Sigma-Aldrich), 10 μM Thioflavin T (Sigma-Aldrich) and 0.1 mg/mL recombinant mouse, human, or bank vole PrP substrate as stated in the figure legends. Octuplicate reactions were seeded each time with 2 μL of serially diluted homogenates, starting from 10^-1^, from CWD-infected mice or bank voles. Tissue homogenates (seeds) were 10-fold serially diluted in RT-QuIC seed dilution buffer (0.05% (w/v) SDS in 1X PBS. The plate was sealed with Nunc Amplification Tape (Nalge Nunc International) and placed in a BMG Labtech FLUOstar Omega fluorescence plate reader that was pre-heated to 42° C for a total of 50 h, or with an additional step of substrate replacement after 25 hours, with cycles of 1-minute double orbital shaking (700 rpm) incubation and 1-minute resting throughout the assay time. For substrate replacement, cycling was stopped after 25 h and 90 μl of the reaction mixture were replaced with fresh assay buffer containing rPrP and cycling was continued up to the end. Thioflavin T fluorescence signals of each well were read and documented every 15 minutes, then the values the relative fluorescence units (RFU) were plotted as the average of octuplicate reactions by using GraphPad Prism (version 9) software. For each assay, corresponding tissue from an age-matched non-infected tg650 or non-infected bank vole was added as internal negative control. The threshold was calculated based on the average fluorescence values of negative control +5SD. Up to three RT-QuIC independent experiments were performed for most of the runs.

### Statistical analysis

GraphPad Prism 9.0 software (GraphPad) was used to establish the heatmap for the RT-QuIC assays. It was also used to draw the RT-QuIC graphs, quantification of glyco-form ratios, and to do the statistical analyses.

## Acknowledgements

We thank Dr. Trent Bollinger, WCVM, University of Saskatchewan, Saskatoon, Canada, for providing brain tissue from the WTD-116AG isolate, Dr. Stéphane Haïk, ICM, Paris, France, for providing brain tissue from vCJD and sCJD cases, and Dr. Umberto Agrimi, Istituto Superiore di Sanità, Italy, for the bank vole model. We thank animal facility staff for animal care, Dr. Stephanie Anderson for veterinary oversight, and Yo-Ching Cheng for preparing recombinant PrP substrates. Thank you to Dr. Stephanie Booth and Jennifer Myskiw, Public Health Agency of Canada, Canada. We are grateful for financial support from NSERC, NIH, Genome Canada, and the Alberta Prion Research Institute. SG is supported by the Canada Research Chairs program.

## Conflict of interests

The authors declare no conflict of interest.

---

**Table S1.** Transmission of CWD prions to tg650 transgenic mice overexpressing human PrP (129MM).

| Animal<br>original<br>ID# | CWD<br>inocula | Days<br>post<br>infection<br>(DPI) | Status of<br>animal at<br>euthanasia | Clinical signs of prion<br>disease | Reason for<br>euthanasia | IHC | RT-QuIC |  |  |  |
| --- | --- | --- | --- | --- | --- | --- | --- | --- | --- | --- |
|  |  |  |  |  |  |  | Brain | Faeces<br>FH dilution: % of positive<br>replicates |  |  |
|  |  |  |  |  |  |  |  | 10<sup>-1</sup> | 10<sup>-2</sup> | 10<sup>-3</sup> |
| #2-HcW-1-321 | Wisc-1 | 882 dpi | Terminal<br>clinical signs | Myoclonus, Weight loss,<br>Hyperexcitability,<br>Kyphosis, Hind-limb<br>claspings, Rigid tail,<br>Rough coat | Terminal | NA | Pos | 12.5% | 12.5% | 12.5% |
| #2-HcW-1-322 |  | 791 dpi | - | No signs of prion disease | Humane<br>endpoint | NA | Pos | Neg | Neg | Neg |
| #2-HcW-1-323 |  | 919 dpi | Terminal clinical signs | Myoclonus, Repeated weight loss and gain, Rigid tail, Kyphosis, Hind-limb claspings | Experimental endpoint | Neg | Pos | 25% | Neg | Neg |
| #2-HcW-1-324 |  | 919 dpi | Subtle clinical signs | Myoclonus, Repeated weight loss and gain | Experimental endpoint | Neg | Pos | Neg | Neg | Neg |
| #2-HcW-1-325 |  | 789 dpi | Terminal clinical signs | Myoclonus, Repeated weight loss and gain, Rigid tail, Kyphosis, Hind-limb claspings, Mild ataxia, Paralysis | Terminal | NA | Pos | Neg | Neg | Neg |
| #2-HcW-1-326 |  | 934 dpi | - | Myoclonus | Experimental endpoint | Neg | Inconclusive | Neg | 25% | 12.5% |
| #2-HcW-1-327 |  | 623 dpi | Terminal clinical signs | Myoclonus, Weight loss, Rigid tail, Kyphosis, Hind-limb claspings, Ataxia, Paralysis, Heavy breathing, Irresponsiveness, Gait abnormalities | Terminal | Neg | Inconclusive/Neg | 75% | 100% | 75% |
| #2-HcW-1-328 |  | 934 dpi | Subtle clinical signs | Myoclonus, Heavy breathing | Experimental endpoint | **Pos** | Pos | Neg | Neg | 12.5% |
| #2-HcW-1-329 |  | 934 dpi | Subtle clinical signs | Myoclonus, Heavy breathing | Experimental endpoint | Neg | Pos | Neg | Neg | Neg |
| #2-HcW-1-330 |  | 213 dpi | - | No signs of prion disease | Found dead | NA | NA | NA | NA | NA |
| #2-HcW-1-331 | 116AG | 882 dpi | Terminal clinical signs | Myoclonus, Weight loss, Hind-limb clasping, Gait abnormalities, Hind-limb paralysis | Terminal | NA | Inconclusive/Neg | 12.5% | 12.5% | 12.5% |
| #2-HcW-1-332 |  | 919 dpi | Subtle clinical signs | Myoclonus, Gait abnormalities | Experimental endpoint | NA | Neg | Neg | Neg | Neg |
| #2-HcW-1-333 |  | 919 dpi | Subtle clinical signs | Myoclonus, Repeated weight loss and gain | Experimental endpoint | NA | Neg | Neg | Neg | Neg |
| #2-HcW-1-334 |  | 722 dpi | - | No signs of prion disease | Humane endpoint | NA | NA | Neg | Neg | Neg |
| #2-HcW-1-335 |  | 751 dpi | - | No signs of prion disease | Found dead | NA | Neg | Neg | 25% | Neg |
| #2-HcW-1-336 |  | 934 dpi | - | Myoclonus | Experimental endpoint | NA | Neg | 12.5% | Neg | 12.5% |
| #2-HcW-1-337 |  | 736 dpi | - | No signs of prion disease | Found dead | NA | Neg | NA | NA | NA |
| #2-HcW-1-338 |  | 919 dpi | Subtle clinical signs | Weight loss, Kyphosis, Gait abnormalities, Myoclonus and hyperexcitable | Experimental endpoint | NA | Neg | 12.5% | 12.5% | Neg |
| #2-HcW-1-339 |  | 934 dpi | Terminal clinical signs | Myoclonus, Kyphosis, Gait abnormalities | Experimental endpoint | NA | Neg | Neg | Neg | Neg |
| #2-HcW-1-340 |  | 934 dpi | - | Myoclonus | Experimental endpoint | NA | Neg | Neg | Neg | Neg |

**Table S2.**
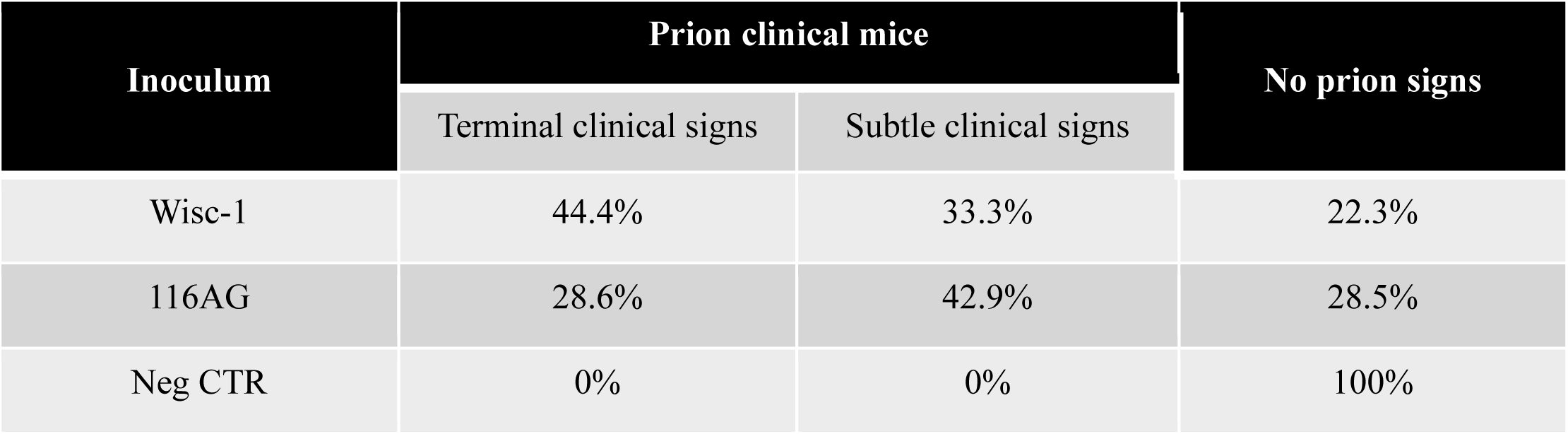
Bioassay of WTD isolates in tg650 mice.

| Inoculum | Prion clinical mice |  | No prion signs |
| --- | --- | --- | --- |
|  | Terminal clinical signs | Subtle clinical signs |  |
| Wisc-1 | 44.4% | 33.3% | 22.3% |
| 116AG | 28.6% | 42.9% | 28.5% |
| Neg CTR | 0% | 0% | 100% |

## Supplementary Data

**Figure S1.**
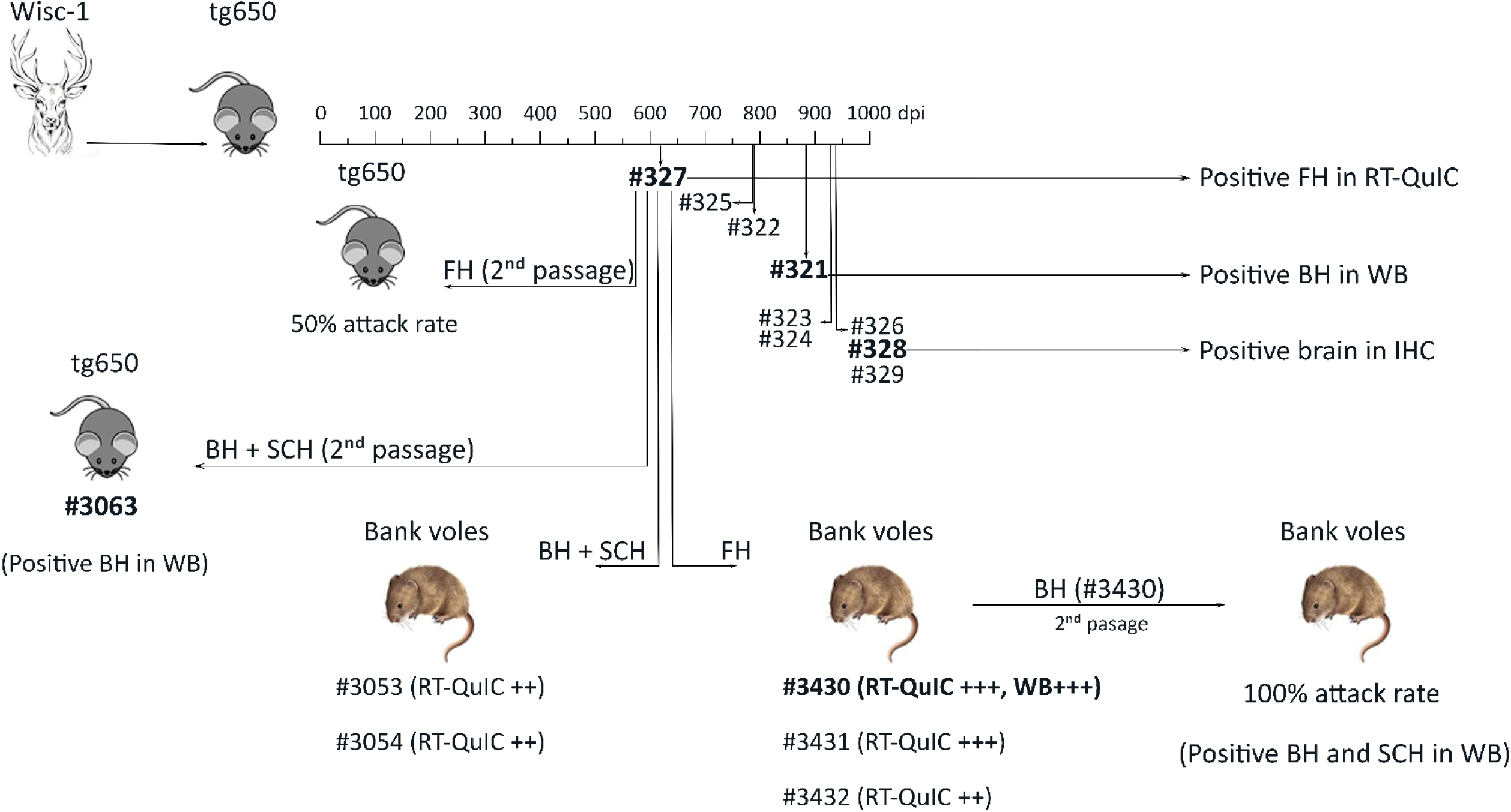
**Scheme of the transmission study of WTD Wisc-1-CWD in tg650 and bank vole models.**

**Figure S2.**
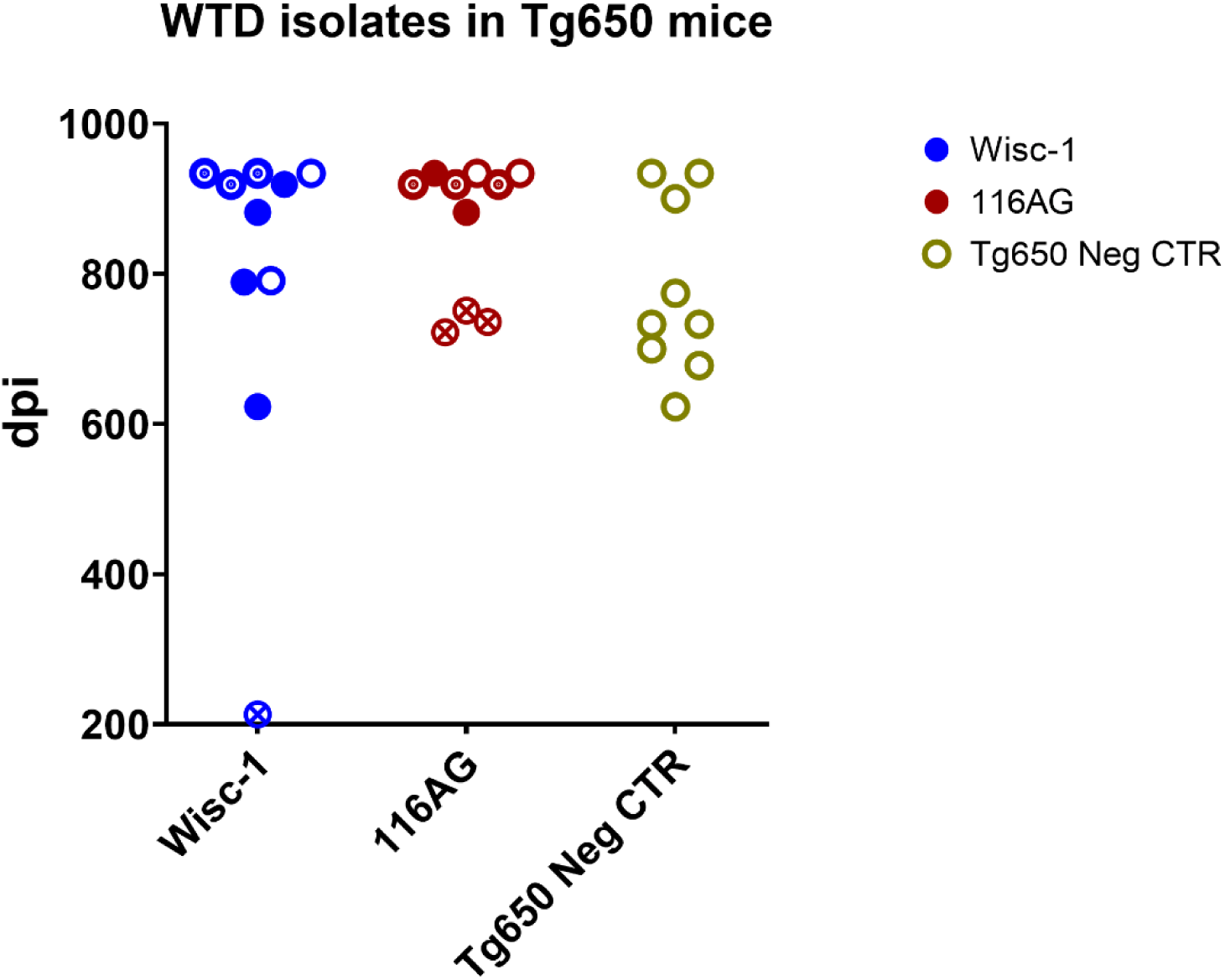
Transmission of WTD isolates in tg650 mice. Tg650 mice were inoculated with Wisc- 1 (blue), 116AG (red), or not inoculated (age-matched control; green). Distribution of incubation period in tg650 mice is classified according to the status of mice at the time of euthanasia. Mice are defined with terminal clinical signs (full circles), subtle clinical (bullet circles), or no signs (open circles). Circles with crosses represent mice euthanized due to intercurrent diseases.

**Figure S3.**
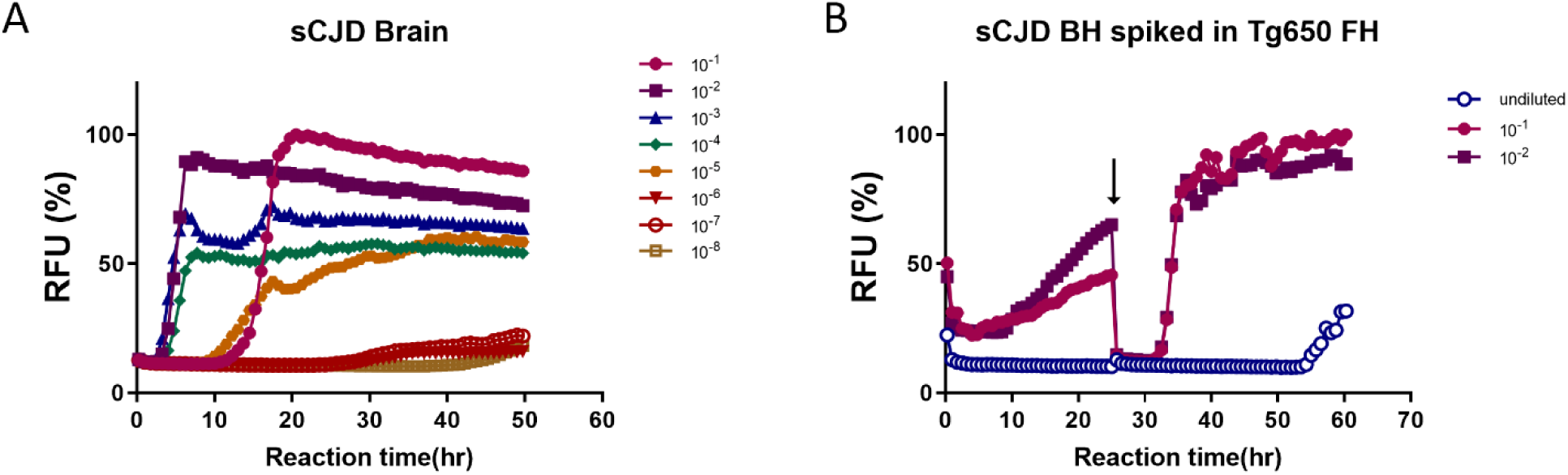
**A.** RT-QuIC analysis of sCJD BH prion seeding activity. The graphs depict a representative RT-QuIC assay of brain homogenates from an sCJD patient-MM1. Twenty percent sCJD brain homogenates were serially diluted (10^-1^ – 10^-8^) and seeded in mouse rPrP substrate. **B.** RT-QuIC analysis of sCJD BH spiked in fecal homogenate of tg650 naïve mouse and seeded in mouse rPrP substrate. Introduction of substrate replacement is indicated with a black arrow.

**Figure S4.**
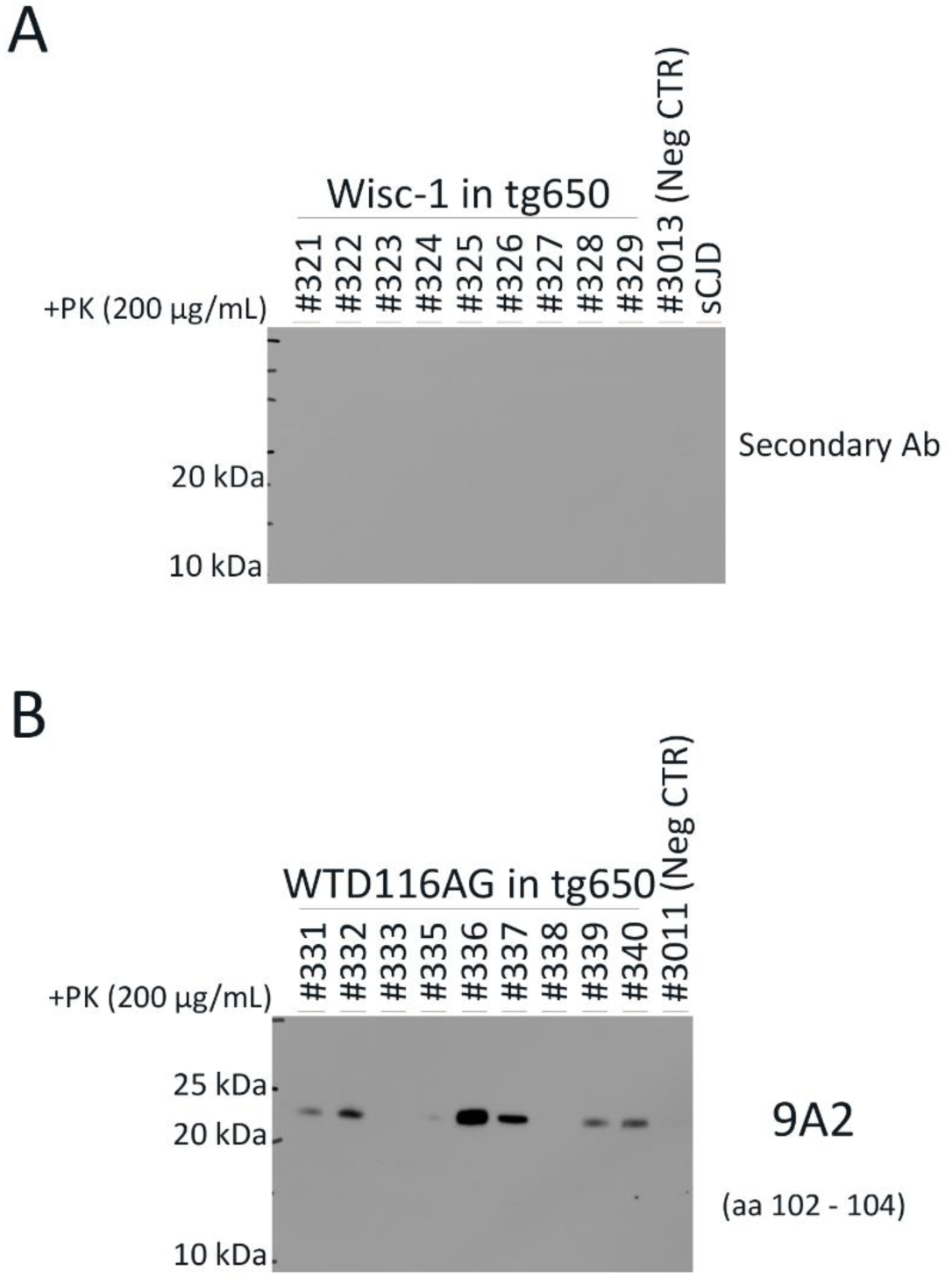
**A.** Western blot analysis of brain homogenates of tg650 mice inoculated with Wisc-1 isolate and digested with 200 µg/mL of PK using only horseradish peroxidase-conjugated goat anti-mouse IgG. **B.** Western blot analysis of brain homogenates of tg650 mice inoculated with 116AG isolate and digested with 200 µg/mL of PK using anti-PrP mAb, 9A2 (aa 102 – 104, bottom). A negative control, tg650 #3011 was also included in the western blot.

**Figure S5.**
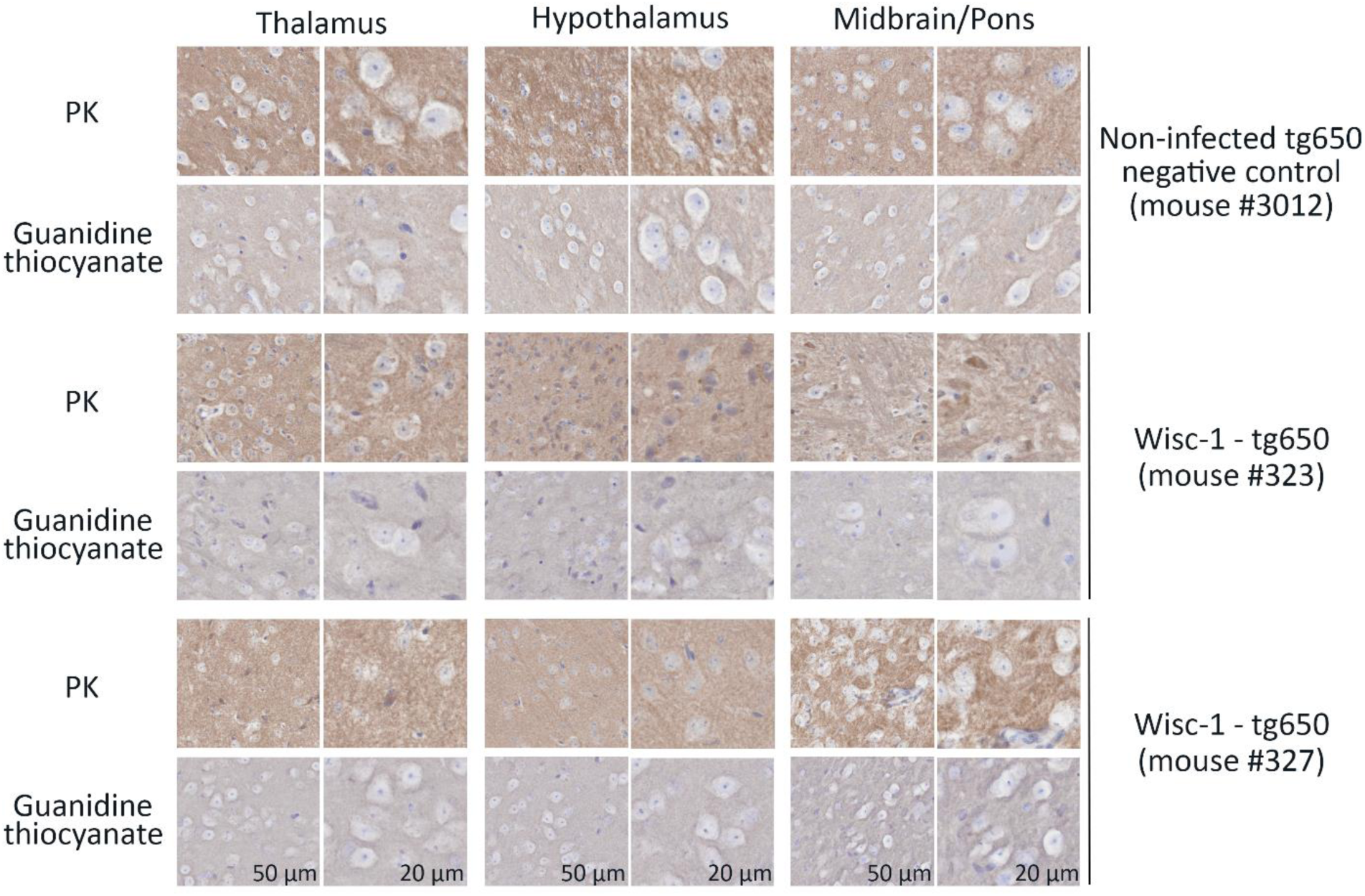
PrP^Sc^ staining of age-matched negative tg650 control, and Wisc-1 inoculated tg650 mice. Immunohistochemistry using either PK digestion (upper panels) or guanidine denaturation (lower panels) demonstrating the lack of PrP^Sc^ staining in the thalamus (left panels), hypothalamus (middle panels), and midbrain/pons (right panels) areas of tg650 negative control #3012 (upper panels), Wisc-1 inoculated tg650 mouse #323 (middle panels), and mouse #327 (lower panels). Scale bars, 50 µm, and 20 µm.

**Figure S6.**
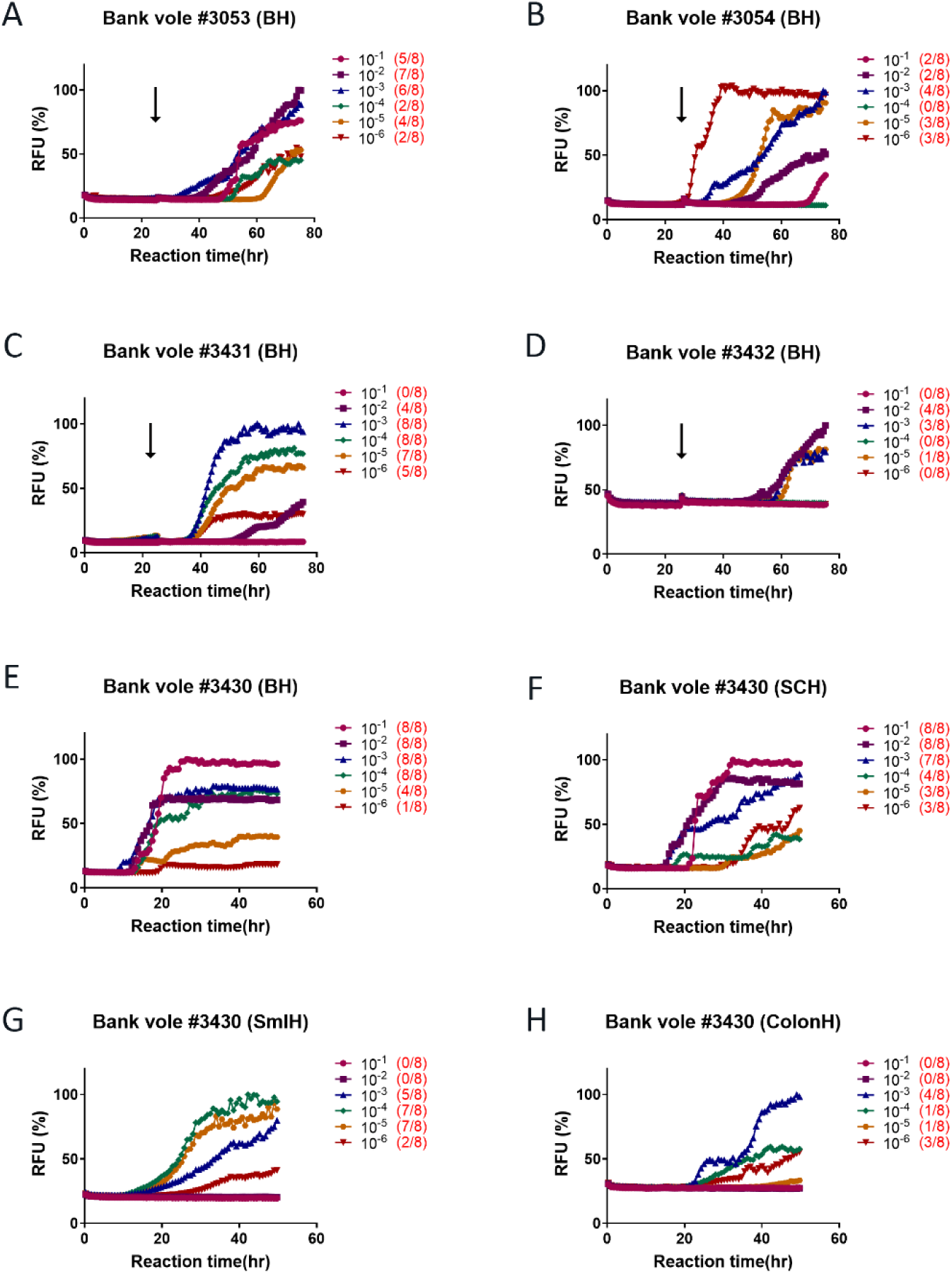
Transmission of CWD-tg650 to bank voles. One percent brain/spinal cord homogenates from Wisc-1-inoculated tg650 mouse was transmitted intracerebrally to bank voles. The curves depict a representative RT-QuIC assay of brain homogenates from bank vole #3053 (A), and #3054 (B). Ten percent fecal homogenates from Wisc-1-inoculated tg650 mouse were inoculated intracerebrally to bank voles. The graphs depict a representative RT-QuIC assay of brain homogenates from bank voles #3431 (C), #3432 (D), and brain (E), spinal cord (F), small intestine (G), and colon (H) homogenates from bank vole #3430. Bank vole rPrP was used as a substrate. The presence of a black arrow in A-D indicates the replacement of substrate with a fresh one.

**Figure S7.**
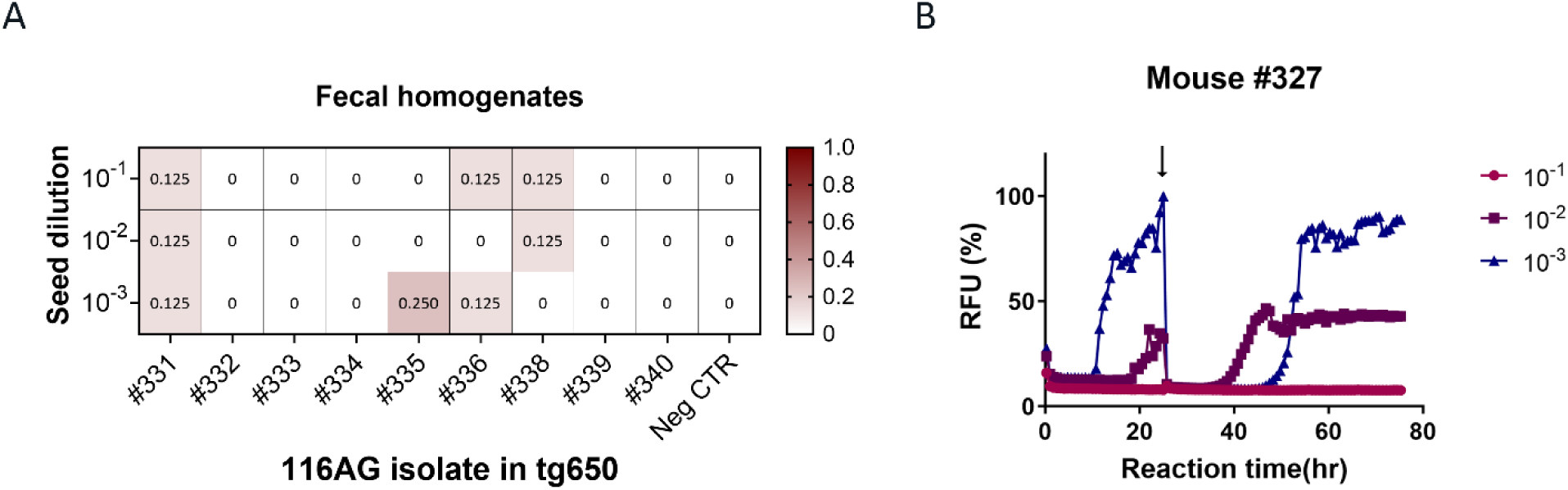
Prion seeding activity detected in feces of 116AG-inoculated tg650 mice. **A.** Summary of RT-QuIC analysis of prion seeding in the fecal material of 116AG humanized inoculated mice. The heatmap indicates the percentage of positive RT-QuIC replicates out of the total of eight replicates analyzed. The scale goes from 0 (all replicates were negative) to 1 (all replicates were positive). Fecal homogenates were serially diluted (10^-1^ to 10^-3^) and mouse rPrP substrate was used as a substrate. **B.** The curves depict a representative RT-QuIC assay of fecal homogenates from Wisc-1-inoculated tg650 mouse #327 serially diluted (10^-1^ to 10^-3^) using human rPrP substrate. The black arrow indicates the substrate replacement.

**Figure S8.**
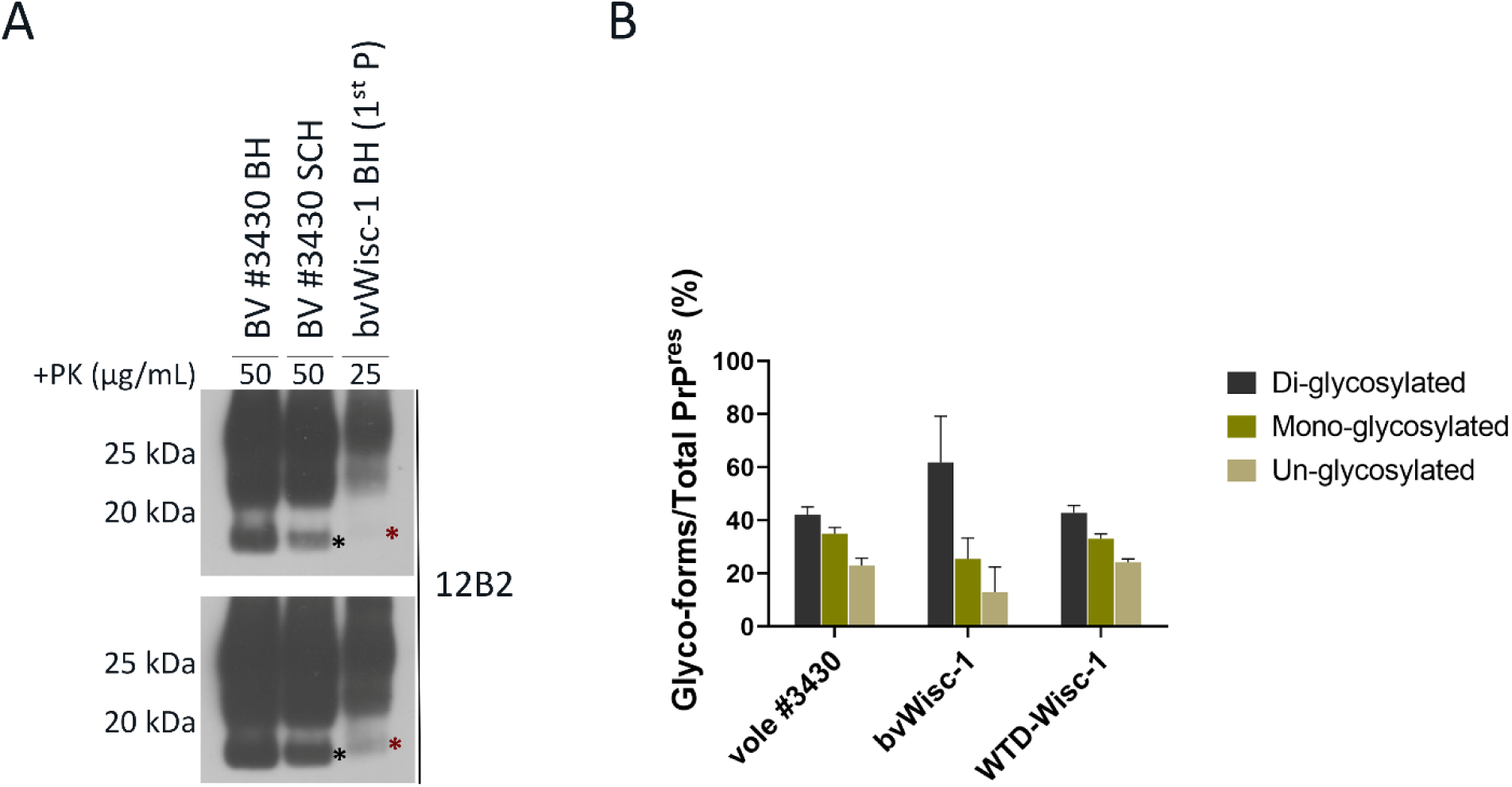
Transmission of CWD-tg650 fecal material to bank voles. 10% fecal homogenate from Wisc-1 inoculated tg650 mouse #327 was inoculated intracerebrally to bank voles. **A.** Western blot analyses of fecal homogenate-inoculated bank vole #3430 using brain homogenates (lane 1) and spinal cord homogenates (lane 2) digested with 50 µg/mL of PK, as well as Wisc-1 passaged in bank voles (bvWisc-1) 1^st^ passage (lane 3) digested with 25 µg/mL of PK. The western blot was probed with mAb 12B2. The amount of bvWisc-1 loaded on the gel was 20x that of bank vole #3430. The upper and lower panels are the same blots with different exposure times to better show the slower migration of the un-glycosylated band depicted in the bvWisc-1 1^st^ passage. **B.** Quantification of the glycoform ratios of Wisc-1 isolate, bank vole #3430, and bvWisc-1 (1^st^ and 2^nd^ passage) prions.

## Notes

### Competing Interest Statement

The authors have declared no competing interest.

